# The genetic basis of replicated bullseye pattern reduction across the Trionum Complex

**DOI:** 10.1101/2024.10.10.617677

**Authors:** May T. S. Yeo, Alice L. M. Fairnie, Valentina Travaglia, Joseph F. Walker, Lucie Riglet, Selin Zeyrek, Edwige Moyroud

## Abstract

Angiosperm flowers exhibit a wide diversity of colorful motifs on their petals. Such patterns fulfill both biotic and abiotic functions, mediating plant-pollinator communication and providing protection against damaging UV rays or desiccation. These motifs are often evolutionary labile, varying in size, shape and hue between closely related species and constitute excellent systems to illuminate the evolutionary processes that generate morphological diversity or instead lead to the repetitive emergence of similar forms. *Hibiscus trionum* flowers have a prominent bullseye pattern combining a purple center contrasting against a white margin. *H. trionum* belongs to a small clade of *Hibiscus* known as the Trionum Complex that displays a range of petal patterns within and between species. Here, we integrate phylogenomic approaches, molecular techniques and genetic manipulation to solve species relationships across the Trionum Complex, identify key genes involved in the production of bullseye pigmentation, and reveal molecular events underpinning pattern variation during the evolution of the group. We find that repetitive bullseye reduction events primarily occur through independent modifications of a single genetic locus encoding BERRY1, an R2R3 MYB that regulates anthocyanin pigment production in petals. Moreover, we demonstrate that buff-tailed bumblebees *(Bombus terrestris)* discriminate against flowers with smaller bullseye sizes suggesting that a reduction in bullseye proportions potentially impacts plant fitness and contributes to trait distribution across the group. Collectively, our results demonstrate how repeated mutations in a single locus led to morphological variation in petal patterning, a trait that contributes to angiosperm reproductive isolation and speciation.

## Introduction

The molecular basis underlying the repeated evolution of similar morphological characteristics across the tree of life remains a central question of evolutionary biology. Replicated evolution, a term encompassing both parallel and convergent evolution, led different species and populations to independently evolve similar features (James *et al*. 2023). While this sometimes relies on the modification of homologous genes (Lenormand *et al*. 2016), it remains unclear why some genes are more likely than others to mediate parallel or convergent changes. The evolution of color patterns in both plants and animals provides phenotypically striking traits that can be utilized to unravel the molecular mechanisms supporting the evolution of diversity (Manceau *et al*. 2010; Orteu & Jiggins, 2020; Galipot *et al*. 2021). With more than 350,000 extant species, the angiosperms are a living repository of natural diversity. The diversification of flowering plants yielded a remarkable array of morphologies and colorful patterns that decorate corollas. Remarkably, similar motifs are often found in distantly related species. Striking examples include petal stripes in *Gazania* and *Phlox*, venation patterns in *Veronica*, *Geranium* and *Antirrhinum*, or bullseye patterns in the poached egg plant (*Limnathes douglasii*) and baby blue eyes (*Nemophila menziesii*). Such commonalities hint towards a limited but functionally important set of patterns that independently arose across distantly related angiosperms. It also provides multiple systems with which to investigate the processes that have led to the repeated emergence of similar traits.

Petal patterns play crucial roles in key biological processes including plant-pollinator communication, defense against herbivory, and protection from abiotic factors and thus are likely to contribute directly to plant fitness (Fairnie *et al*. 2022). Insect pollinators can display innate or acquired preferences for given motifs (Gumbert, 2000), influencing their choice of flower to visit. Importantly, petal patterns also function post-landing, acting as visual guides to lure pollinators toward pollen and/or nectar rewards and increase the chances of successful pollination. The lower petal lobes of the purple monkeyflower (*Mimulus lewisii*), for example, display prominent yellow markings surrounded by dark pink stripes with a white border. Mutants lacking these nectar guides or with reduced petal coloration exhibit significant reductions in pollinator visitation (Owen & Bradshaw, 2011). Further work in yellow monkeyflowers (*Mimulus guttatus*) reinforced the importance of pigmentation patterns for pollinator attraction as bumblebees preferred the flowers of the *red tongue* mutant with an expanded anthocyanin region over wildtype flowers with smaller anthocyanin spots (Ding *et al*. 2020). Petal patterns also play important roles to protect against abiotic stress. In wild sunflowers (*Helinathus annuus*), larger UV-absorbing bullseyes increased a flower’s tolerance to desiccation (Todesco *et al*. 2022). In silverweed flowers the size of the UV bullseye positively correlates with the amount of exposure to UV irradiation, possibly enhancing pollen protection from UV-induced DNA damage (Koski & Ashman, 2015). In both species, the UV-absorbing bullseye results from the preferential or restricted production of flavonols at the petal base which protect against environmental stress (Pollastri & Tattini, 2011). Hence, the evolution of petal patterns is shaped by both biotic and abiotic factors and investigating the processes mediating changes in petal patterning provides insights into the mechanisms facilitating the emergence of adaptations and speciation.

The patterns on the petal surface are produced by a combination of features including epidermal cells that differ in their shape, texture, and pigmentation (Kay, 1981; Johnson & Midgley, 1997). Plant pigments are divided into three classes: carotenoids, betalains and flavonoids. The carotenoid and flavonoid biosynthetic pathways are widely distributed across the plant kingdom while betalains are restricted to the order Caryophyllales. Flavonoids are the predominant class of pigments in angiosperm flowers and produce a wide array of colors.

Flavonoids comprise several subgroups, including flavonols and anthocyanins, which are colorless/‘cream’-like and blue/red, respectively. Flavonoid biosynthesis begins with the production of chalcone, followed by several hydroxylation steps to produce the precursors for flavonols and anthocyanins. Thereafter, the pathway branches whereby the enzymes dihydroflavonol (*DFR*) and anthocyanidin synthase (*ANS*) produce anthocyanins while flavonol synthase (*FLS*) yield flavonols, respectively (Tanaka *et al*. 2008). The accumulation of these pigments, in varying combinations and abundance, and in different regions of the petals, gives rise to the spectrum of color patterns on the petal epidermis. Petal coloration can also be influenced by the physical properties of epidermal cells (Moyroud *et al*. 2017; Riglet *et al*. 2021). For instance, conical cells can focus light into the anthocyanin-containing vacuole enhancing color intensity, while semi-ordered striations of the cuticle can produce a weak iridescence and blue halo effect that increases flower salience to pollinators (Whitney *et al*. 2009; Moyroud *et al*. 2017). The genetic components underpinning the production of petal patterns have been functionally characterized in a limited number of model species. Flavonoid biosynthesis is regulated at the transcriptional level by the MBW complex, combining two types of transcription factors (TFs), an R2R3 MYB (myeloblastosis) and a basic helix-loop-helix (bHLH) protein, and a scaffold WD-repeat protein (Ramsey & Glover, 2005). MYBs and bHLHs constitute two of the largest TF families in plants and as such, several MBW complexes exist to tightly regulate the expression of late flavonoid biosynthetic genes. In *Antirrhinum majus,* the venation pattern present on the lateral sides of the corolla tube is produced by overlapping expression patterns of the MYB TF AmVENOSA and its bHLH partner, AmDELILA. Together, they combine with a WD40 co-activator, forming an MBW complex that activate anthocyanin biosynthesis (Shang *<u>et al.</u>* 201<u>1</u>). *AmVENOSA* expression is restricted to a wedge of cells between the vein and adaxial epidermis while *AmDELILA* is broadly expressed in the epidermal cells of the corolla (Goodrich *et al*.1992; Jackson *et al*. 1992). Therefore, the restricted expression of *AmVENOSA*, the MYB element of the MBW complex, is the main driver of the venation pattern. The mechanisms accounting for the spatial regulation of *MYB* transcription has only been explored in a few systems including *Mimulus verbenaceus* leaf stripes (LaFountain *et al*. 2024) because many species used in the study of petal patterning cannot be genetically manipulated with ease. Hence, the upstream processes accounting for the restriction of MYB expression are still poorly understood across angiosperms.

Venice mallow (*Hibiscus trionum)* produces flowers with a striking bullseye pattern, and this herbaceous species has recently emerged as a model system to understand petal pattern evolution and development at both the macro- and microscale (Moyroud *et al*. 2022; Riglet *et al*. 2024). *H. trionum* belongs to the Trionum clade within the Malvaceae family and specifically falls into the Trionum Complex that appears to have experienced a recent radiation event (Craven *et al*. 2011). The size of the bullseye varies extensively across the Trionum Complex providing an attractive system to investigate the processes underpinning petal pattern variation and possibly replicated evolution. In this study, we first used a phylogenomic approach to solve species relationships across the Trionum Complex and uncovered multiple instances of bullseye pigmentation reduction. Combining molecular approaches and genetic manipulations *in planta*, we found that independent occurrences of bullseye size reductions are due to recurring modifications of a MYB-encoding locus, *BERRY1*. This locus restricts anthocyanin production to the petal base and acts as a principal driver of bullseye pattern formation. Importantly, pollinator behavior assays revealed that bumblebees easily discriminate between flowers of varying bullseye size, suggesting that repeated mutations within a single locus are sufficient to generate morphological variations in a trait likely to impact speciation.

## Results and Discussion

### Bullseye pattern reduction occurred several times independently across the Trionum Complex

To investigate petal pattern evolution across the Trionum Complex, we first undertook a phylogenomic approach to clarify the relationships between the four species within the group (Fig. 1A). Whenever possible, we included several accessions from each species to capture some of the morphological diversity existing within some species (Fig. 1B) (Craven *et al*. 2011; Moyroud *et al*. 2022). For instance, *H. verdcourtii* individuals from the Emerald and Theodore localities in Queensland (referred to as population 1 and 4, respectively) exhibit a red bullseye, while New South Wales individuals from the Narrabri region (population 5) produce cream flowers lacking a pigmented bullseye (Fig. 1B). Individuals with different bullseye patterns can also be found in the same region. For example, plants producing red bullseye flowers (*H. verdcourtii* population 2) or cream flowers with a faint pink halo (*H. verdcourtii* population 3) are both present in St. George, Queensland (Craven *et al*. 2011, Fig. 1B). Similarly, a commercial accession of *H. trionum* used in previous studies (van der Kooi *et al*. 2015; Moyroud *et al*. 2022) produces a visibly smaller bullseye than the one of *H. trionum* CUBG, our wildtype reference accession (Moyroud *et al*. 2022; Riglet *et al*. 2024) and wild individuals from New Zealand (*H. trionum* Bream Head, Craven *et al*. 2011; Moyroud *et al*. 2022) (Fig. 1B).

**Fig. 1:**
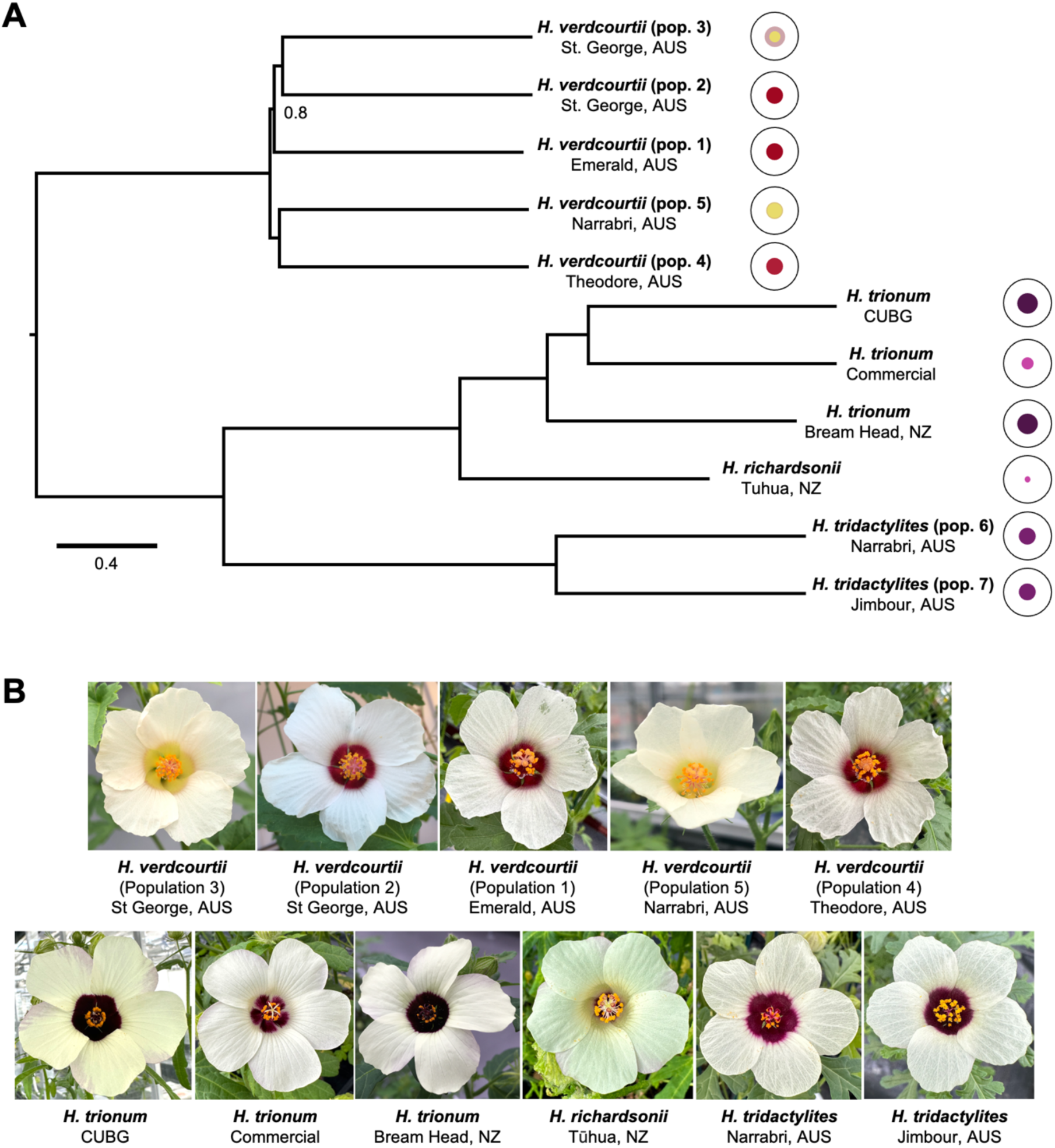
Phylogenetic relationships between Trionum Complex species. **(A)** Phylogenetic tree highlighting relationships between species and populations. Unless stated, all nodes are perfectly supported. Branch lengths are shown in coalescent units. Each species is represented by a cartoon depicting bullseye size and color. **(B)** Representative flowers of each species/populations emphasizing variation in bullseye size and color.

As expected, accessions assigned to the same species clustered together in our phylogenomic reconstruction (Fig. 1A). Our analysis also identified *H. richardsonii*, the other diploid species of the group (de Lange, 2008; Murray *et al*. 2008), as sister to *H. trionum*. This finding is in line with previous morphological and ecological studies where the status of *H. richardsonii* as a separate species or a variant of *H. trionum* was debated (de Lange, 2008; Craven *et al*. 2011). We found that all accessions produced smooth conical cells in the distal part of the petal and striated tabular cells in the proximal petal region except for the five *H. verdcourtii* accessions that all produced smooth tabular cells in the proximal petal region, regardless of their corolla pigmentation pattern (Fig. S1). This indicates (i) that the genetic mechanisms controlling cell shape, cuticle texture, and pigment production are somewhat distinct and (ii) that the ability to produce a striated cuticle was either gained after the divergence of *H. verdcourtii* or lost along the lineage leading to *H. verdcourtii*. Finally, our phylogenetic reconstruction identified pigmentation as the most labile feature of epidermal petal cells as accessions lacking pigmentation at the petal base (*H. verdcourtii* populations 3 and 5) or producing a reduced pigmented bullseye (*H. trionum* Commercial Variety and *H. richardsonii*) did not cluster together (Fig. 1A). Therefore, understanding the genetic basis accounting for multiples instances of bullseye pigmentation reduction provides an excellent opportunity to study the mechanisms mediating replicated evolution.

### A single genetic locus accounts for reduction of the pigmented bullseye in *H. richardsonii*, the sister species of *H. trionum*

We previously found that the boundary separating proximal and distal regions is specified closer to the petal base during the pre-patterning phase in *H. richardsonii* compared to wildtype *H. trionum* (Riglet *et al*. 2024). This boundary shift is sufficient to produce a smaller structural bullseye in *H. richardsonii* as striated tabular cells cover a smaller portion of its petal. However, the pigmented area appears even smaller than the proximal domain suggesting that additional changes affecting the control of pigment production, rather than the pre-patterning process itself, must also have occurred during evolution along the lineage leading to *H. richardsonii* (Riglet *et al*. 2024). Here, we aim to clarify the genetic basis accounting for such reduction in pigmentation. To precisely characterize differences in pigmentation between wildtype *H. trionum* and *H. richardsonii* bullseyes, we first examined and compared the morphological and chemical features of fully open flowers (stage 5) from both species. We found that the pigmented bullseye covered on average 15% of the petal surface in wildtype *H. trionum* while the *H. richardsonii* bullseye represented only 3% of the petal surface (Fig. 2B). This five-fold reduction in pigmented bullseye size was mainly attributed to a drastic reduction in anthocyanin levels in the *H. richardsonii* proximal region compared to wildtype *H. trionum*, with anthocyanin content of the proximal region being at least 15-fold lower in *H. richardsonii* compared to *H. trionum* (Fig. 2C). Minor pigment accumulation in the distal petal <u>is</u> likely attributed to light-induced anthocyanin production on the abaxial petal. Total flavonol content was 1.4 to 2.6-fold higher in *H. richardsonii* petal tissue compared to wildtype *H. trionum* (Fig. 2D). Taken together our data indicate that the pigmented bullseye of *H. trionum* is generated by restricting anthocyanin synthesis to the proximal petal region while flavonols are preferentially produced in the distal petal domain and that bullseye reduction in *H. richardsonii* is likely due to impaired anthocyanin synthesis. Similarly, in the *Clarkia gracilis* species complex, petal spots vary among subspecies and the presence of basal and central petal spots was correlated to higher anthocyanidin levels in those regions compared to unspotted petals that had lowered anthocyanidin levels (Martins *et al*. 2013). In *H*. *richardsonii*, the reduction in anthocyanin production could favor precursor availability toward the flavonol branch of the flavonoid pathway, possibly accounting for the increase in flavonols recorded in the proximal and boundary regions of *H. richardsonii* petals (Fig. 2D).

**Fig. 2:**
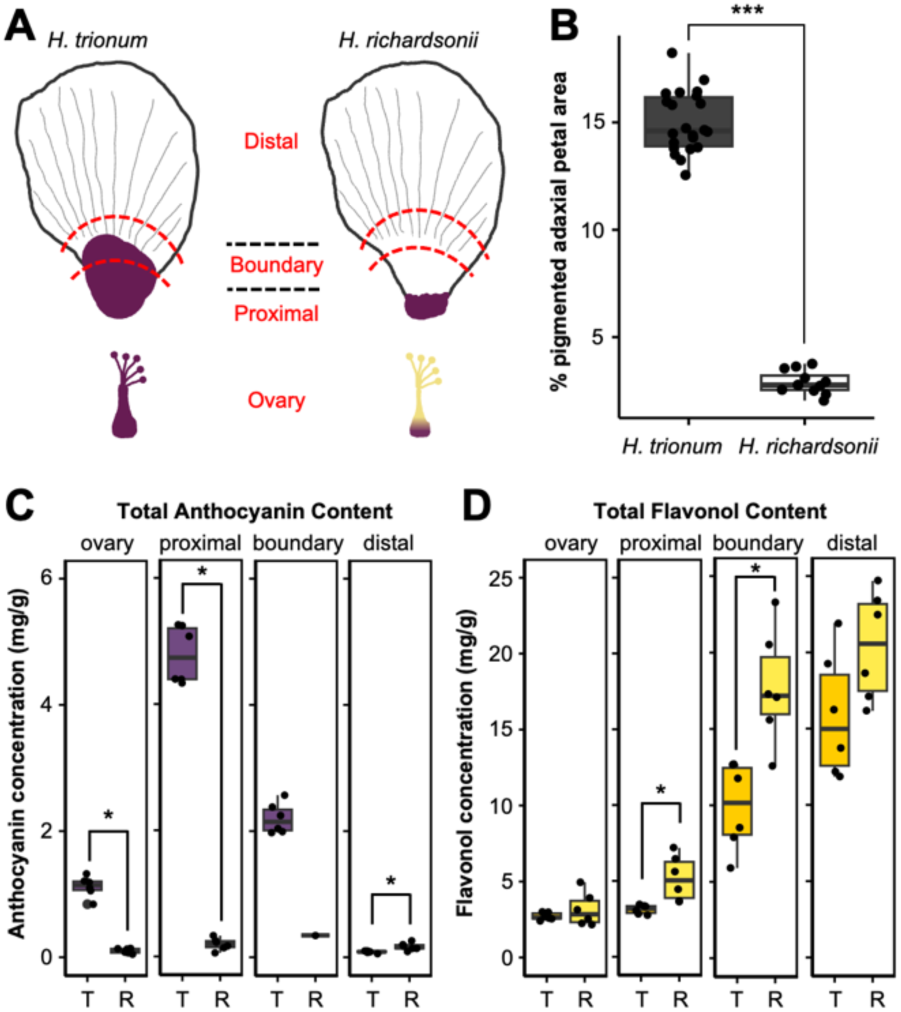
Morphological differences in petal patterning between wildtype *H. trionum* and *H. richardsonii.* **(A)** Schematic depicting petal regions and ovary of a stage 5 flower. For pigment extraction assays, petals were dissected into ovary, proximal, boundary and distal regions. Each region is demarcated by the red dashed lines. **(B)** The size of the pigmented bullseye was measured as a percentage of the pigmented region over total petal area. 22 and 12 flowers were measured from wildtype *H. trionum* and *H. richardsonii,* respectively. 3 petals were measured per flower. *** p<0.001 (Student’s t-test). **(C)** Anthocyanin and **(D)** flavonol quantification of wildtype *H. trionum* (T) and *H. richardsonii* (R) flowers. Stage 5 flowers were dissected into 4 regions as highlighted in (A). 6 flowers were analyzed per species. Each data point represents the average of three technical replicates. * p<0.05 (Welch’s or Student’s t-test) of pairwise comparisons made between species in the same region.

To start uncovering the molecular mechanisms accounting for the difference in bullseye pigmentation between wildtype *H. trionum* and *H. richardsonii,* we crossed the two species together (Fig. 3A). The hybrid produced flowers with a pigmented bullseye similar in size to wildtype *H. trionum* (Fig. 3A and 3B). Closer inspection of the boundary region (Fig. 3C) revealed differences in the composition and pattern of cell types between the hybrid and wildtype *H. trionum* parent. In the parental wildtype *H. trionum*, the boundary region was characterized by three cell types: (1) striated pigmented tabular cells, (2) smooth pigmented tabular cells, and (3) smooth non-pigmented tabular cells (Fig. 3C). In *H. richardsonii,* the reduction in bullseye pigmentation resulted in only two cell types at the boundary: (1) striated and (2) smooth non-pigmented tabular cells. Interestingly in the hybrid, striated non-pigmented tabular cells replaced the smooth pigmented tabular cells found in wildtype *H. trionum*. This indicates that although the hybrid appears identical to wild type *H. trionum* at the macroscale, the dominance of the *H. trionum* phenotype is somewhat partial as the hybrid only displays a near full-size pigmented proximal region. In the F2 population obtained after the hybrid was allowed to self-pollinate, the two parental and hybrid phenotypes were recapitulated and these morphologies segregated in a near-Mendelian ratio as 40, 41 and 96 individuals displayed a *H. trionum*-like, *H. richardsonii*-like and hybrid-like phenotype, respectively. This indicates the change in pigmented bullseye dimensions is largely due to a single locus (Fig. 3). Similarly, a single Mendelian locus controls the presence of “light areas” on the proximal petal in *M. lewisii.* The hybrid from a cross between *M. lewisii* and *M. cardinalis* displayed the light areas phenotype, indicating the *M. lewisii* allele is dominant for this pigment pattern trait (Yuan, *et al*. 2016).

**Fig. 3.**
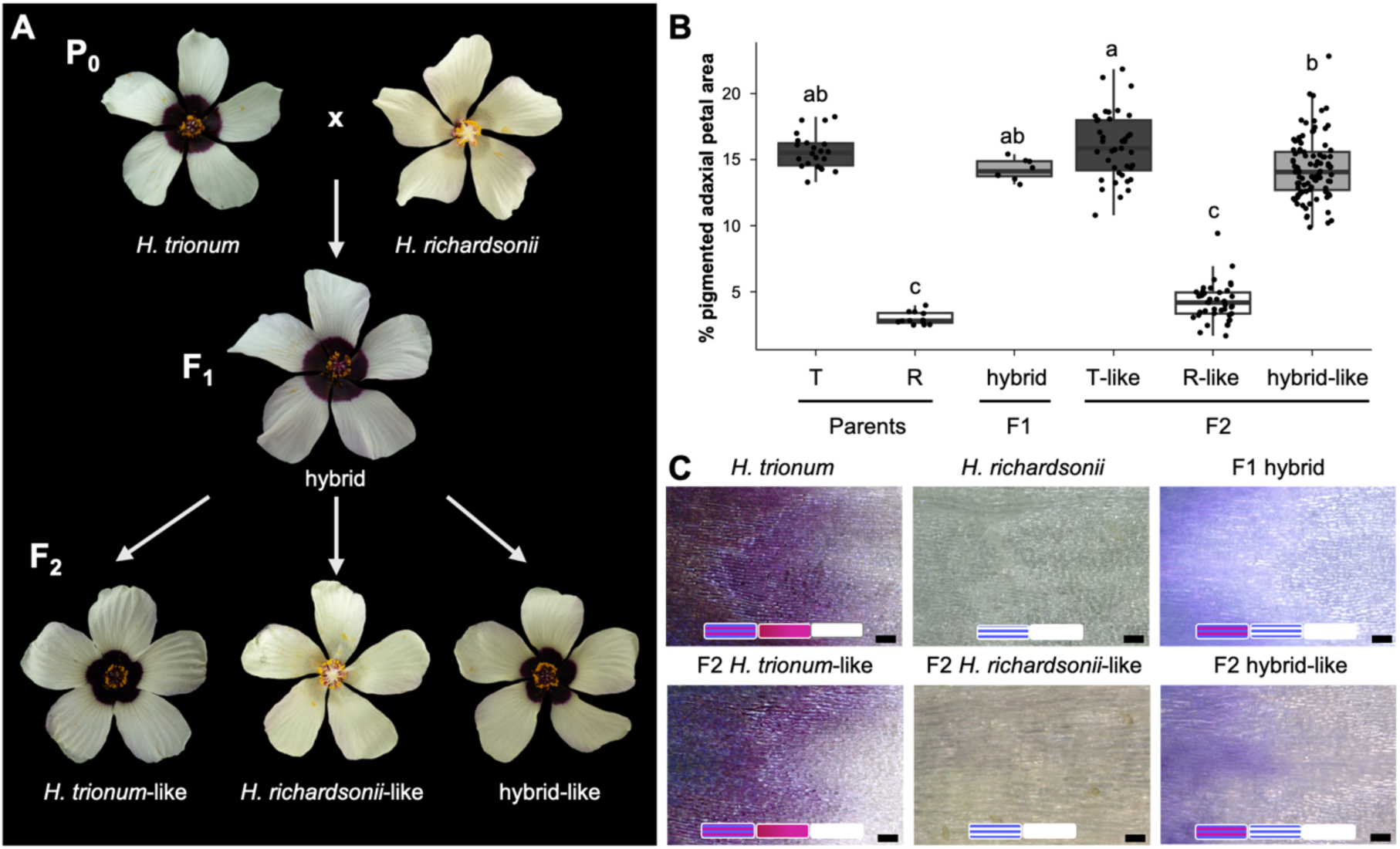
Characterization of wildtype *H. trionum, H. richardsonii,* F1 and F2 phenotypes. **(A)** F1 and F2 flower phenotypes of *H. trionum* crossed with H. richardsonii. The F1 hybrid produced a flower with a pigmented bullseye. The F2 selfing population yielded three flower phenotypes that recapitulate the two parental and F1 hybrid flower morphologies. **(B)** Measurement of the pigmented bullseye size reflects F2 phenotypes that match parental (*H. trionum* = T, *H. richardsonii* = R) and F1 hybrid phenotypes. Different letters indicate statistically significant differences (Tukey’s HSD, p < 0.0004). **(C)** Imaging of the boundary region of petals from the six flower morphologies from (A). Cell shape (tabular) is consistent between all samples. Moving left to right along the proximo-distal axis: in the *H. trionum* wildtype and F2 *H. trionum*-like boundaries, cells transitioned from striated to non-striated and lost pigmentation. In the H. richardsonii and F2 *H. richardsonii*-like boundaries, pigmentation was absent, and striation was lost. In the F1 and F2 hybrid-like boundaries, both pigmented and non-pigmented striated cells were present, with striation loss occurring after pigmentation loss. Scale bar = 100um.

### Expression patterns of a *DFR* and a *FLS Hibiscus* homolog correlate with petal pigmentation

The flavonoid pathway is often targeted by evolutionary processes, generating diversity across many flowering species (Kellenberger & Glover, 2023). Therefore, we hypothesized that differences in petal patterning between *H. trionum* and *H. richardsonii* is also a consequence of modifications to this pathway. The flavonoid biosynthetic pathway is well-characterized in plants (Fig. 4A). A common dihydroflavonol precursor is produced before the pathway diverges to synthesize anthocyanins or flavonols. *DFR* and *FLS* are key enzymes involved in the production of anthocyanins and flavonols, respectively. From the *H. trionum* genome, we isolated three homologs for each gene. We measured their expression as petals developed (stages 1-4, prior to flower opening, Moyroud *et al*. 2022) in the proximal and distal regions (Figs. 4B and 4C). Among the three *DFR* homologs, *HtDFR1* was expressed at the highest levels in the proximal region: *HtDFR1* expression was eight to 11-fold higher at the petal base compared to the distal region throughout petal development (Figs. 4B). *HtDFR2* and *HtDFR3* expression levels in the proximal petal were almost negligeable and thus we concluded that they are not likely involved in anthocyanin production at the petal base (Fig. S2 and S3). *HtFLS2* was highly expressed during the first three stages of petal development (Fig. 4C) before declining at stage 4, following a similar temporal dynamic yet complementary spatial expression pattern to *HtDFR1* as its expression was consistently four to 19-fold higher in the distal region (Fig. 4C, Fig. S3). Neither *HtFLS1* nor *HtFLS3* were expressed at levels comparable to *HtFLS2* (Figs. S2 and S3). Taken together, these results demonstrate that *HtDFR1* and *HtFLS2* are the main players in shaping the bullseye pattern, and the spatial accumulation of anthocyanins and flavonols (Fig. 2C-D) directly reflects their expression patterns. In *Dianthus caryophyllus, Rosa multiflora,* and *Camellia japonica* flowers, expression of *DFR* and *FLS* homologs also correlate with red and white flowers, respectively (Luo *et al*. 2016). In *Clarkia gracilis, DFR* expression patterns coincide with petal spots formation during development (Martins *et al*. 2013). In our findings, we also demonstrate compartment-specific expression of *DFR* and *FLS* prior to the emergence of pigmentation patterns at very early stages of development (stage 1, Figs. 4 and Fig. S3). Our results also argue for functional divergence between the *DFR* and *FLS* homologs. Such diversification involved divergence of their expression patterns but whether this was accompanied by changes in coding sequences affecting the biochemical properties of the enzymes they code for remains to be determined.

**Fig. 4.**
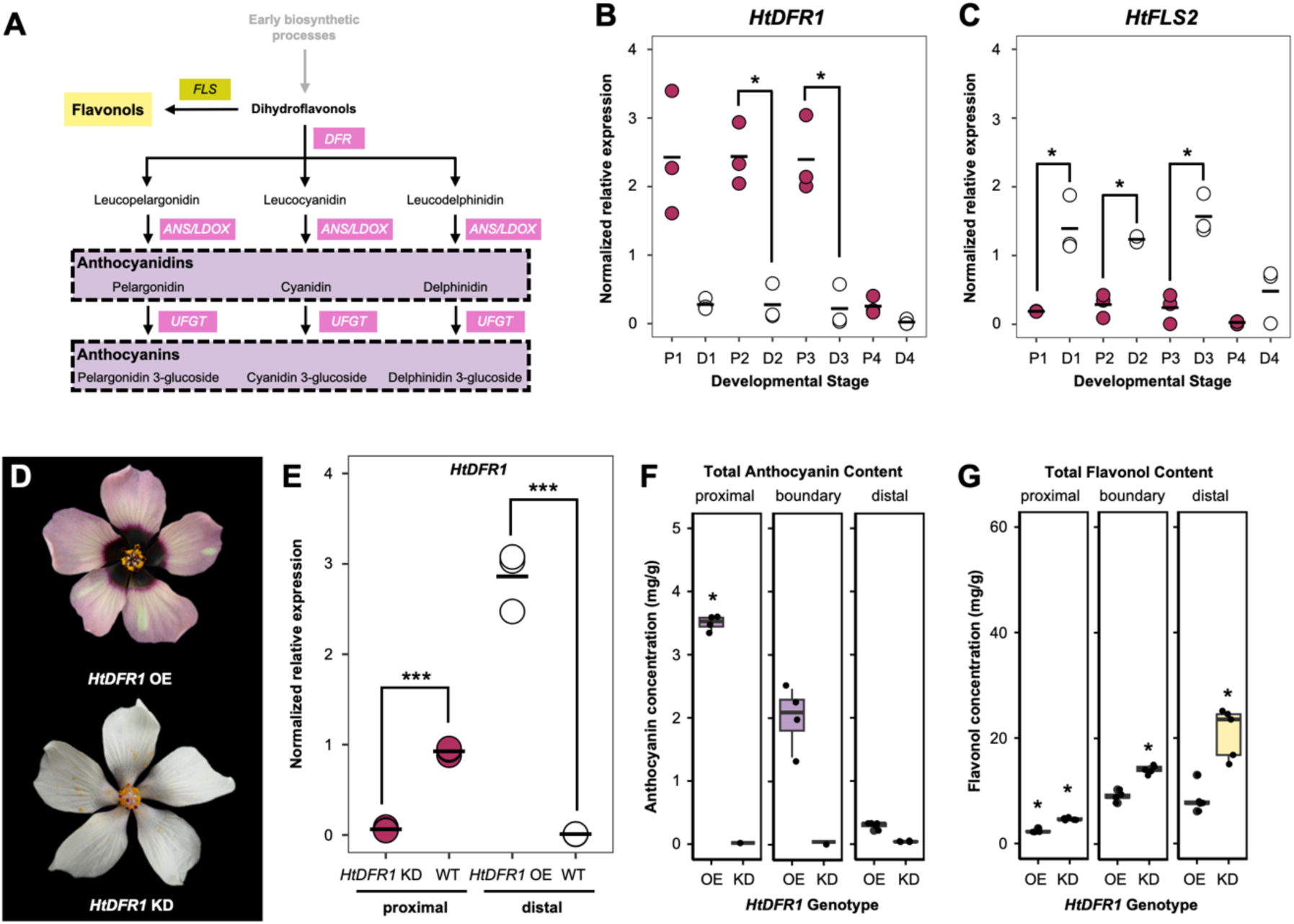
Bullseye pigmentation reflects patterning of *DFR* and *FLS* expression across the petal primordia. **(A)** Simplified flavonoid biosynthesis pathway highlighting dihydroflavonols as a shared common precursor for flavonol and anthocyanin production via *FLS* and *DFR* enzyme activity, respectively. Expression of **(B)** *HtDFR1* and **(C)** *HtFLS2* throughout development in proximal (P1 to P4) and distal (D1 to D4) petal tissue of wildtype *H. trionum.* * denotes p<0.05 (Student’s or Welch’s t-test) of pairwise comparisons made between proximal and distal tissue of the same stage. **(D)** Phenotypes of two *35S::HtDFR1* lines. **(E)** Quantification of *HtDFR1* expression in *35S::HtDFR1* KD and OE lines. *** denotes p<0.01 (Student’s or Welch’s t-test) for comparisons of *HtDFR1* against WT in the same petal region. **(F)** Anthocyanin and **(G)** flavonol quantification in both *35S::HtDFR1* lines. Stage 5 petals were dissected into 3 regions as highlighted in Fig. 2A. 5 flowers were analyzed per line. Each data point represents the average of three technical replicates. * denotes p<0.05 (Student’s, Wilcox’s or Welch’s t-test) of pairwise comparisons made between *HtDFR1* OE vs. WT and *HtDFR1* KD vs. WT tissue of the same region. Comparisons were not made in regions that had fewer than three data points as some replicates yielded non-detectable amounts of anthocyanin. For expression data in (B), (C) and (E), three biological replicates were extracted per stage and each data point indicates an average of three technical replicates, horizontal bars indicate mean relative expression values.

To determine whether *HtDFR1* was responsible for bullseye anthocyanin production, we generated *35S::HtDFR1* transgenic lines in *H. trionum*. Strong overexpression lines with ectopic pigmentation in the distal petal were observed, however, a single transgenic line produced flowers devoid of a pigmented bullseye (Fig. 4D). We hypothesized that this was likely due to silencing of *HtDFR1* triggered by transcript overaccumulation similar to results in *Arabidopsis* T-DNA supertransformants (Schubert *et al*. 2004). To test this hypothesis, we quantified *HtDFR1* transcription levels in transgenic lines exhibiting either phenotype (Fig. 4D). We found that the presence of distal pigmentation was accompanied with a 300-fold averaged increase in *HtDFR1* expression in the distal region (Fig. 4E). Reciprocally, *HtDFR1* transcription was minimal (15-fold lower than WT expression) in the proximal petal domain of white flowers, confirming that *HtDFR1* expression matched the strength of the phenotype observed. We concluded that the absence of pigmentation was likely due to *HtDFR1* silencing, such that the transgenic line producing white flowers corresponded to a knockdown (KD) line for *HtDFR1*. Total anthocyanin and flavonol measurements in both lines confirmed visual observations: anthocyanin levels were five-fold higher in the distal petal region of the *HtDFR1* overexpression line (Fig. 4F). Conversely, flavonol levels were two-fold higher throughout the entire petal of the *HtDFR1* knockdown line (Fig. 4G). The lack of *HtDFR1* activity in the knockdown line likely increased precursor availability for flavonol biosynthesis, possibly accounting for increased flavonol levels.

Although *HtDFR1* overexpression produced distal petal pigmentation, anthocyanin levels in the distal petal domain remained 13-fold lower compared to anthocyanin production in the proximal region suggesting that other elements, such as substrate competition between ectopic HtDFR1 and endogenous HtFLS2, could limit anthocyanin production. A clear bullseye pattern was still visible in the *HtDFR1* knockdown line despite the near-complete absence of anthocyanin (Fig. 4D), highlighting the contribution of the structural elements (differences in cell shape and texture between the proximal and distal region) to the visual appearance of the petal. This also indicates that the genetic changes accounting for pigment reduction in *H. richardsonii* must only affect the gene regulatory network regulating pigment production instead of the upstream processes initiating boundary formation during the pre-patterning phase (Riglet *et al*. 2024).

To identify the genetic changes accounting for reduced anthocyanin production in *H. richardsonii*, we isolated *HrDFR1* and *HrFLS2* in *H. richardsonii* and did not find any key differences in amino acid sequences when compared against their orthologs in wildtype *H. trionum*. Next, we measured their expression throughout *H. richardsonii* petal development and found that compared to wildtype *H. trionum*, *HrDFR1* expression was significantly reduced across all petal stages in both proximal (17- to 107-fold lower) and distal (two- to nine-fold lower) tissue (Fig. 5A). The expression profile of *HrFLS2* was similar to *HtFLS2* such that its expression was also higher in the distal petal compared to the proximal petal. Hence, the reduction of bullseye pigmentation in *H. richardsonii* correlates with a severe reduction in *HrDFR1* expression across the petal primordia.

**Fig. 5.**
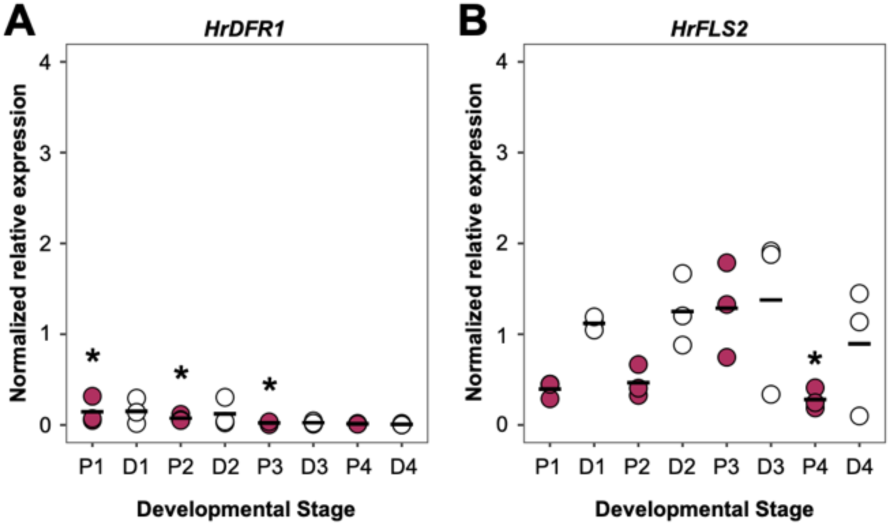
(A) Expression of *HrDFR1* and **(B)** *HrFLS2* throughout development in proximal (P1 to P4) and distal (D1 to D4) petal tissue of H. richardsonii. (Fig. 4B). Three biological replicates were extracted per stage. Each data point indicates an average of three technical replicates, horizontal bars indicate mean relative expression values. * denotes p<0.05 (Student’s t-test) of pairwise comparisons made between the same region of wildtype *H. trionum* and *H. richardsonii*.

### Changes at the *BERRY1* locus are responsible for differences in pigmented bullseye size between *H. trionum* and *H. richardsonii*

The regulation of genes encoding key enzymes such as *DFR* and *FLS* in the flavonoid biosynthesis pathway is in part controlled by MBW complexes, comprising a MYB, a bHLH and a WD40 component (Xu *et al*. 2014). We reasoned that some of the TFs regulating *HtDFR1* transcription must be preferentially expressed in the proximal region of the petal. To identify such regulatory genes, we dissected stage 1 petal primordia lacking a visible pigmented bullseye into distal and proximal regions and performed RNAseq to detect differentially expressed genes (Fig. S4A, Supplementary Data 1 & 2). Using a four-fold difference as a threshold, we identified 938 and 343 genes preferentially expressed in the proximal or distal petal regions, respectively (Fig. S4B). Among those ∼1300 genes, three MYBs stood out: two that were almost exclusively expressed in the proximal petal domain, which we named *HtBERRY1* and *HtBERRY2*, and one, that we named *HtCREAM1*, was preferentially expressed in the distal petal region. A phylogenetic analysis of the MYB family revealed that the two *HtBERRY* genes belong to subgroup 6, a clade that comprises the *A. thaliana PAP1/2* genes as well as several known regulators of pigment production in snapdragon, petunia, monkeyflowers and orchids (Fig. S5A, Supplementary Data 3 & 4). *HtCREAM1* fell within a clade comprising cocoa and cotton representatives but lacking *Arabidopsis* genes (Fig. S5B), making it difficult to predict its possible role.

qRT-PCR analysis in wildtype *H. trionum* confirmed the differential expression of *HtBERRY1/2* and *HtCREAM1* along the proximo-distal axis of the petal throughout development. *HtBERRY1* transcription was always much higher (at least 15-fold) in proximal tissue, increasing in expression until stage 3 before reducing to almost negligible levels (Fig. 6A). *HtBERRY2* expression mirrored that of *HtBERRY1* but its overall transcription level was lower compared to its paralog. *HtCREAM1* expression complemented *HtBERRY1/2* expression patterns such that its expression was always higher in the distal tissue at early stages (at least five-fold higher in the distal region) but dropped off by stage 3 (Fig. 6A).

**Fig. 6.**
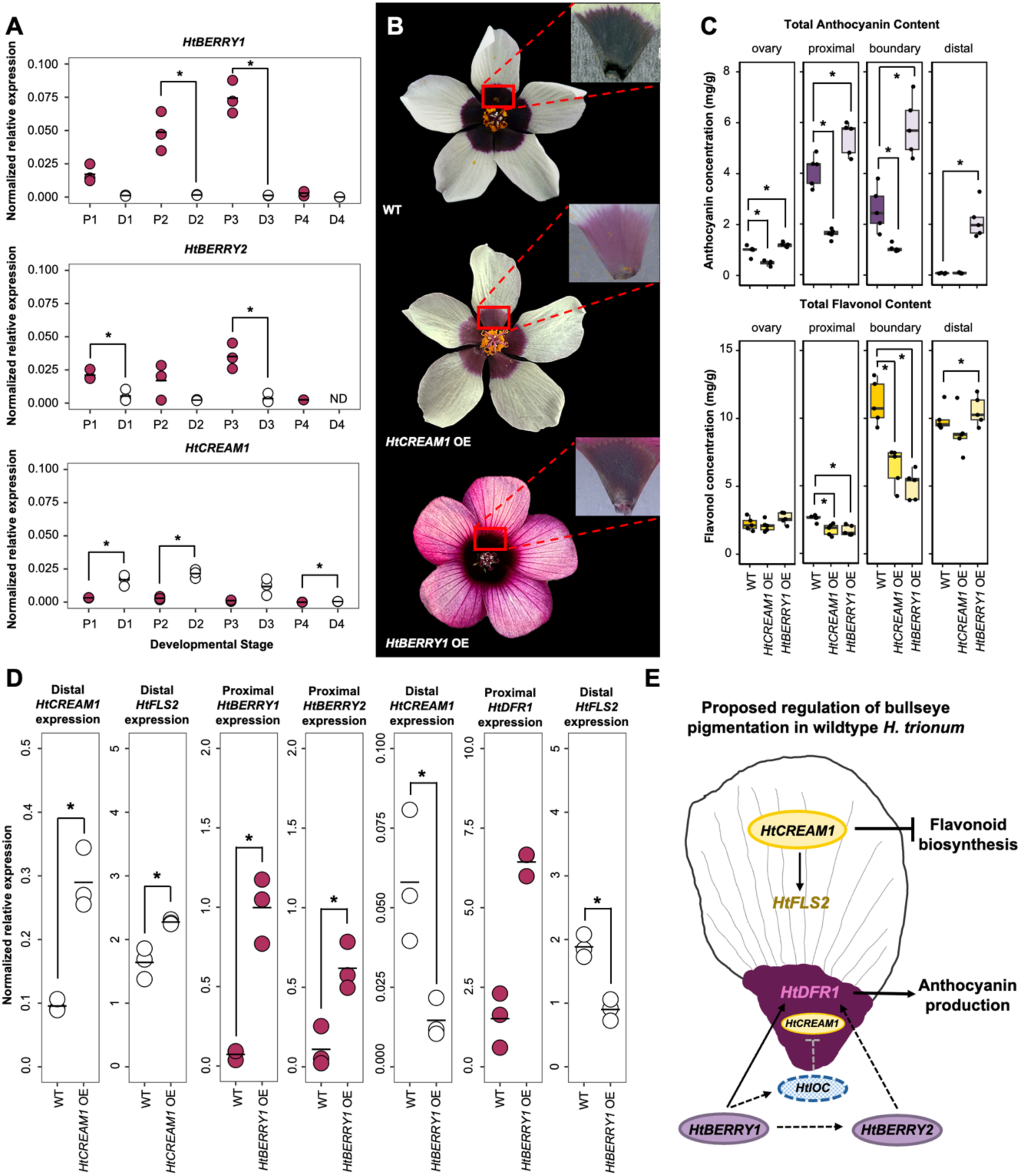
Bullseye pigmentation reflects patterning of *HtBERRY1/2 and HtCREAM1* across the petal primordia. **(A)** Expression of *HtBERRY1*, *HtBERRY2* and *HtCREAM1* MYB TFs throughout development in proximal (P1 to P4) and distal (D1 to D4) petal tissue of wildtype *H. trionum*. ND = non-detectable expression. * denotes p<0.05 (Student’s or Welch’s t-test) pairwise comparisons made between proximal and distal tissue of the same stage. **(B)** Overexpression phenotypes of 35S::*HtBERRY1 and* 35S::*HtCREAM1.* **(C)** Anthocyanin and flavonol quantification in 35S::*HtBERRY1 and* 35S::*HtCREAM1* lines. Stage 5 flowers were dissected into 4 regions as highlighted in Fig. 2A. 5 flowers were analyzed per line. Each data point represents the average of three technical replicates. **(D)** Expression of flavonoid-related structural and MYB transcription factors in distal or proximal regions of the overexpression lines. * denotes p<0.05 (Student’s or Welch’s t-test) of pairwise comparisons made between the same region against WT and the transgenic line in (C) and (D). **(E)** Proposed regulation of bullseye pigmentation in wildtype *H. trionum*. IOC = inhibitor of CREAM1. For expression data in (A) and (D), three biological replicates were extracted per stage and each data point indicates an average of three technical replicates, horizontal bars indicate mean relative expression values.

To investigate the possible functions of these three *MYB* genes and establish their contribution, if any, to bullseye pigmentation, we produced overexpression lines of *HtBERRY1* and *HtCREAM1*. Based on a high degree of sequence similarity (82.9% identity and 94.8% similarity), and overlapping patterns of expression between *HtBERRY1* and *HtBERRY2*, we assumed both paralogs are likely to act redundantly. Therefore, we focused on *HtBERRY1* for functional investigations as it was expressed at higher levels than *HtBERRY2* during petal development. The *35S::HtBERRY1* line (*HtBERRY1* OE) produced flowers with ectopic pink pigmentation in the distal petal: their corolla was entirely colored as the distal region produced 30-fold more anthocyanin than in wildtype (Fig. 6C), however a distinct bullseye pattern was still visible (Fig. 6B). This phenotype was consistent with the predicted role of *HBERRY1* as a positive regulator of anthocyanin production. We also measured higher anthocyanin levels in the ovary and all other petal regions of this line (Fig. 6C), corroborating the ectopic pigmentation and the ability of HtBERRY1 to promote anthocyanin production. Flavonols levels decreased by 30% and 50% in the proximal and boundary regions, respectively, when compared to wildtype *H. trionum*. This could be due to competition between anthocyanin and flavonol production with the increase in anthocyanin biosynthesis resulting in less precursor available for flavonol synthesis. Flowers from the *35S::HtCREAM1* (*HtCREAM1* OE) line were more similar to wildtype but displayed a paler bullseye (Fig. 6B). Flavonoid quantification indicated this was due to reduced production of anthocyanin pigments (60% decrease compared to wildtype *H. trionum*) in the proximal region (Fig 6C). Interestingly, the lack of anthocyanin was not accompanied by increased flavonol synthesis: we found that flavonol content was reduced to levels resembling that of the *HtBERRY1* OE line (Fig. 6C), suggesting that when constitutively overexpressed, *HtCREAM1* could dampen the whole flavonoid pathway.

To begin constructing the gene regulatory network controlling pigment production in the *H. trionum* petal, we measured the expression of the three MYB regulators along with the structural genes *HtDFR1* and *HtFLS2* in both *HtBERRY1* and *HtCREAM1* OE lines (Fig. 6D). We found increased *HtFLS2* expression in the distal region of *HtCREAM1* OE petal (1.5-fold) suggesting HtCREAM1 promotes *HtFLS2* transcription. In the *HtBERRY1* OE line, *HtBERRY1*, *HtBERRY2* and *HtDFR1* expression in the proximal petal increased by ∼15-, six- and four-fold compared to wildtype *H. trionum*, respectively. These results indicate *HtBERRY1* promotes anthocyanin production in part through the activation of *HtDFR1* transcription. Given the similarity in coding sequences between the two paralogs, we assume that HtBERRY2 is also likely to activate *HtDFR1* transcription. Our results support the existence of a positive feedback loop where *HtBERRY1* promotes its own expression as well as *HtBERRY2* transcription. Interestingly, both *HtCREAM1* and *HtFLS2* expression decreased (four- and 1.7-fold, respectively) in the distal petal when *HtBERRY1* was constitutively overexpressed. This suggests that HtBERRY1 could repress the expression of *HtCREAM1* and perhaps *HtFLS2* directly or indirectly (via *HtCREAM1*). Taken together, our results allow us to start building a small gene regulatory network accounting for the restriction of anthocyanin production to the proximal petal region (Fig. 6E).

The bullseye pattern is a consequence of the spatial restriction of *HtDFR1* activity to the petal base. This spatially restricted expression is due, in part to the localized expression of one of its activators, *HtBERRY1,* to the proximal petal region as constitutive overexpression of *HtBERRY1* was sufficient to trigger ectopic expression of *HtDFR1* and yield anthocyanin production in the distal region. Interestingly, the coloration obtained in the distal domain in *HtBERRY1* OE line was not as intense as in the proximal domain and the anthocyanin levels remained 12-fold lower (Fig. 4D-F). This could be due to several reasons. First, a lack of or restricted availability of a HtBERRY1 partner could limit its ability to activate *HtDFR1* expression in the distal region (alternatively, other TFs contributing to *HtDFR1* expression might be lacking in the distal region). Secondly, differences in the chromatin landscape at the *HtDFR1* promoter could impair effective binding of HtBERRY1 to *cis*-regulatory elements. However, our expression data (Figs. 6D and S6) suggests that *HtDFR1* is expressed at similar levels in both proximal and distal regions in the *HtBERRY1* OE line and rules out both hypotheses. Alternatively, competition with *HtFLS2* for precursor availability could prevent *HtDFR1* from synthesizing anthocyanin effectively in the distal petal region, or limited expression of enzymes acting downstream of *HtDFR1* in that same region could account for dampened anthocyanin production. The fact that *HtCREAM1* and *HtFLS2* are both downregulated in the distal region of *HtBERRY1* OE make it unlikely that competition between *HtDFR1* and *HtFLS2* is the main cause of the paler hue of the distal domain of *HtBERRY1* OE line. Taken together, our results suggest that HtBERRY1 promotes pigment production by at least three different routes: first, by inducing the expression of *HtDFR1*; second, by promoting its own expression as well as the expression of its paralog *HtBERRY2*; and third, by reducing the transcription of *HtCREAM1* and *HtFLS2*, limiting possible competition between *HtDFR1* and *HtFLS2* in the proximal region and thus favoring anthocyanin synthesis over flavonol production in the bullseye center. Whether HtBERRY1 directly represses *HtCREAM1* and/or *HtFLS2* transcription remains to be tested – if this is the case, HtBERRY1 must be able to act as both an activator and a repressor by forming different MBW complexes involving different bHLH partners. The R2R3 MYB subfamily, to which HtBERRY1 belongs, is the largest of the four MYB families and can be characterized by two adjacent imperfect MYB repeats (Wu *et al*. 2022). Aside from other functions, more than half of R2R3 MYBs are involved in the regulation of specialized metabolism such as phenylpropanoid biosynthesis which result in end products such as flavonoids. The importance of MYB regulation in the flavonoid pathway has been demonstrated numerous times in several species including *Petunia hybrida* and *Antirrhinum majus.* PhDPL, PhPHZ, AmROSEA1/2 and AmVENOSA all induce expression of *DFR* orthologs (Schwinn *et al*. 2006; Albert *et al*. 2011; Shang *et al*. 2011). However, a dual role for an R2R3 MYB as transcriptional activator and repressor has not been previously reported.

Having identified some of the genetic interactions regulating pigment production in the petal of *H. trionum* (Fig. 6K), we hypothesized that one of those interactions or actors of this network was possibly impaired in *H. richardsonii*, interfering with anthocyanin biosynthesis (Fig. 2C). First, the expression of *HrCREAM1* could be enhanced in *H. richardsonii*, repressing anthocyanin production. Alternatively, the expression of the *BERRY* genes could be reduced, yielding a smaller pigmented bullseye in *H. richardsonii*. To test these hypotheses, we isolated orthologs of *BERRY1/2* and *CREAM1* in *H. richardsonii* and measured their expression in developing petals of *H. richardsonii* (Fig. 7A). The expression of *HrCREAM1* was three- to 76-fold lower, not higher, than that of *HtCREAM1* in both proximal and distal petal regions, allowing us to reject the first hypothesis (Fig. 7A). *HrBERRY1* expression was barely detectible and *HrBERRY2* transcription was also significantly reduced by at least 2.5-fold in the proximal region, likely accounting for the lowered expression of *HrDFR1* in developing petals of *H. richardsonii* (Fig. 5A).

**Fig. 7.**
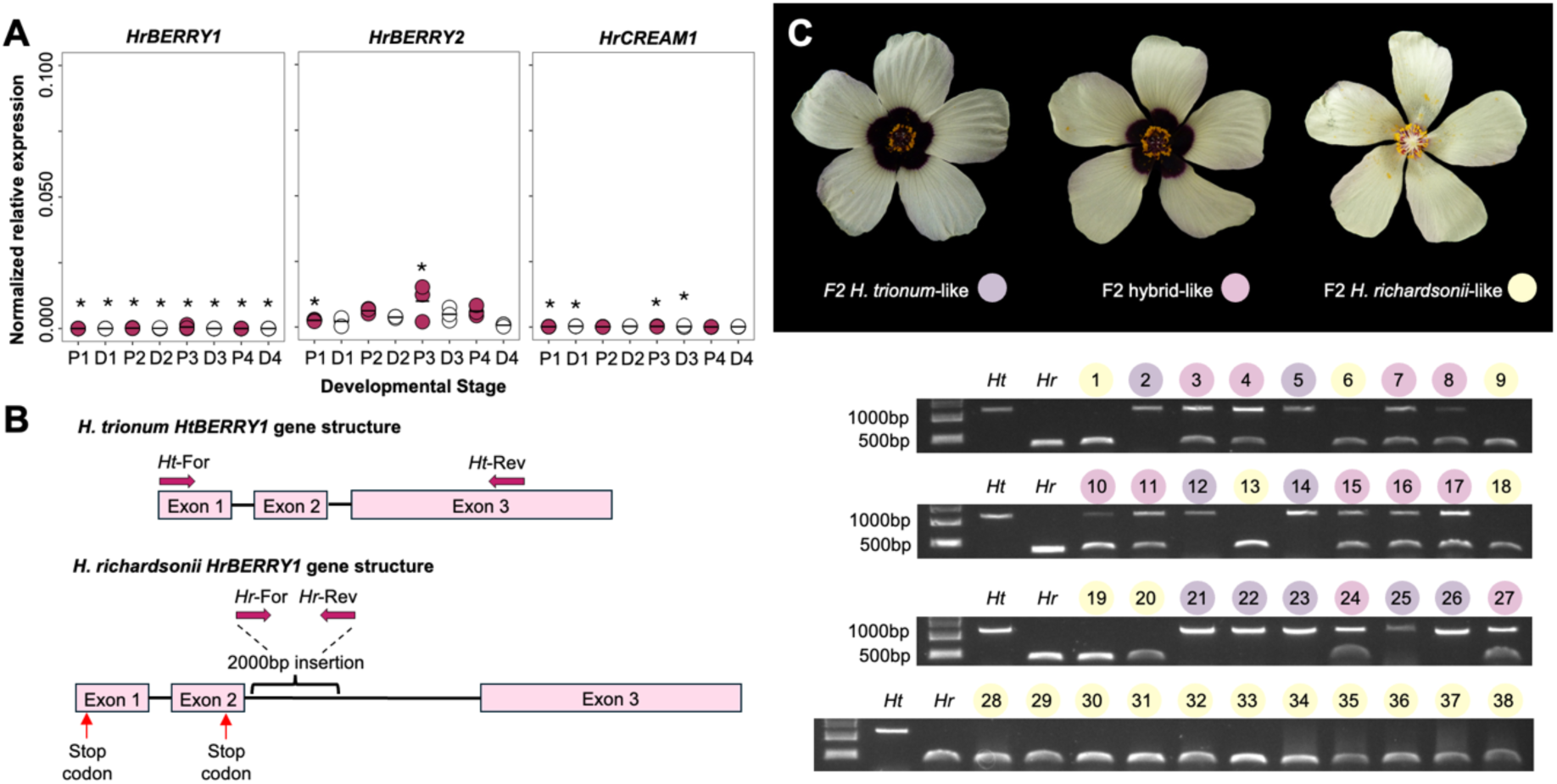
Changes at the *HrBERRY1* locus is responsible for bullseye reduction in *H. richardsonii*. **(A)** Expression of *HrBERRY1, HrBERRY2* and *HrCREAM1* throughout development in proximal (P1 to P4) and distal (D1 to D4) petal tissue of *H. richardsonii*. * denotes p<0.05 (Student’s or Welch’s t-test) of pairwise comparisons made between the same region of wildtype *H. trionum* and *H. richardsonii*. Three biological replicates were extracted per stage and each data point indicates an average of three technical replicates, horizontal bars indicate mean relative expression values. **(B)** Sequencing of *HrBERRY1* revealed two premature stop codons in exons one and two and a 2000bp insertion in intron two. Allele-specific primers used in Fig. 7C are are indicated above each gene structure. **(C)** Genotyping of the *H. trionum (Ht)* x *H. richardsonii (Hr)* F2 population. Blind genotyping of 27 F2 individuals using allele-specific primers revealed that the non-functional *HrBERRY1* allele segregates with the *H. richardsonii-like* F2 phenotype. An additional 11 F2 individuals with the *H. richardsonii*-like phenotype were included to prove segregation.

We also examined the coding sequence of each MYB regulator. We identified a single base pair substitution (A>T, 105bp) in the first exon of *HrCREAM1* but this was a synonymous mutation encoding a glycine residue at position 35. We found the coding sequence of *HrBERRY1* contained eleven SNPs compared to its orthologue in *H. trionum* (three substitutions in exon one, four in exon two, and four in exon three) and a large 2063bp insertion in the second intron (2445bp instead of the 382bp-long intron found in the second intron of *wildtype H. trionum*). Two of the SNPs within the exons introduced premature stop codons: a T>A substitution at position 17bp and G>A substitution at position 159bp. Thus, the *HrBERRY1* transcript should produce short, truncated proteins that are likely non-functional (Fig. 7B). Taken together, our data suggest a scenario where the low expression of *HrDFR1* is caused by a lack of HrBERRY1 activity. If this is the case, the functional *H. trionum* and non-functional *H. richardsonii BERRY1* alleles should segregate with the bullseye size phenotype. To test if the presence of a non-functional *HrBERRY1* allele correlated with a reduced bullseye phenotype, we examined the segregation of the *HtBERRY1* and *HrBERRY1* alleles in the F2 population first described in Fig. 3A. In a blind experiment, 38 F2 individuals were genotyped using allele-specific primers highlighted in Fig. 7B. We found that in all 38 F2 individuals, the genotype at the *BERRY1* locus matched the F2 phenotype (Fig. 7C): all *H. trionum*-like F2 plants were homozygous for the *HtBERRY1* allele, all*. richardsonii*-like individuals were homozygous for the *HrBERRY1* allele, and all hybrid-like plants carried a copy of each allele. Hence, we concluded that the reduction in bullseye pigmentation was likely due to the presence of a non-functional *BERRY1* gene in *H. richardsonii*.

### A loss of BERRY1 activity also accounts for absence of bullseye pigmentation in some populations of *H. verdcourtii*

Bullseye size reduction occurred several times across the Trionum Complex, including within *H. verdcourtii*, generating interspecific variation (Fig. 1B). Hence, changes in bullseye features could precede speciation. Interestingly, we could amplify a *BERRY1* ortholog, *HvBERRY1*, in populations 1, 2 and 4 who all yield individuals with red bullseye flowers, but we failed to amplify this gene in populations 3 and 5 whose flowers are devoid of red pigmentation in the proximal petal region (Fig. 1B). To determine if *BERRY1* was involved in *H. verdcourtii* pigmentation variation, we crossed red bullseye individuals from population 2 with plants from population 3 lacking a pigmented bullseye (Fig. 8A). The F1 hybrid produced flowers resembling those from population 2 (Fig. 8A), suggesting that the pigmented bullseye trait is dominant in *H. verdcourtii*. The hybrid was then allowed to self-pollinate to generate an F2 population. We found that in the F2 population, bullseye color segregated in a 3:1 ratio as 37 and 11 individuals displayed a red and pale bullseye phenotype, respectively. Similar to the findings in the F2 population of wildtype *H. trionum* x *H. richardsonii*, pigmentation loss was largely due to a change affecting a single locus in *H. verdcourtii*. Blind genotyping of 34 F2 individuals revealed that the presence/absence of *HvBERRY1* was a perfect predictor of the presence/absence of a red bullseye (Fig. 8B), supporting the idea that the loss of *HvBERRY1* accounts for the loss of petal pigmentation in *H. verdcourtii*. Taken together, our results reveal a case of replicated evolution, where the pigmented bullseye underwent several independent reductions across the Trionum Complex. These modifications were caused by independent genetic changes (reduction in gene expression, premature stop codons, and gene loss) affecting the *BERRY1* locus (Fig. 8C).

**Fig. 8.**
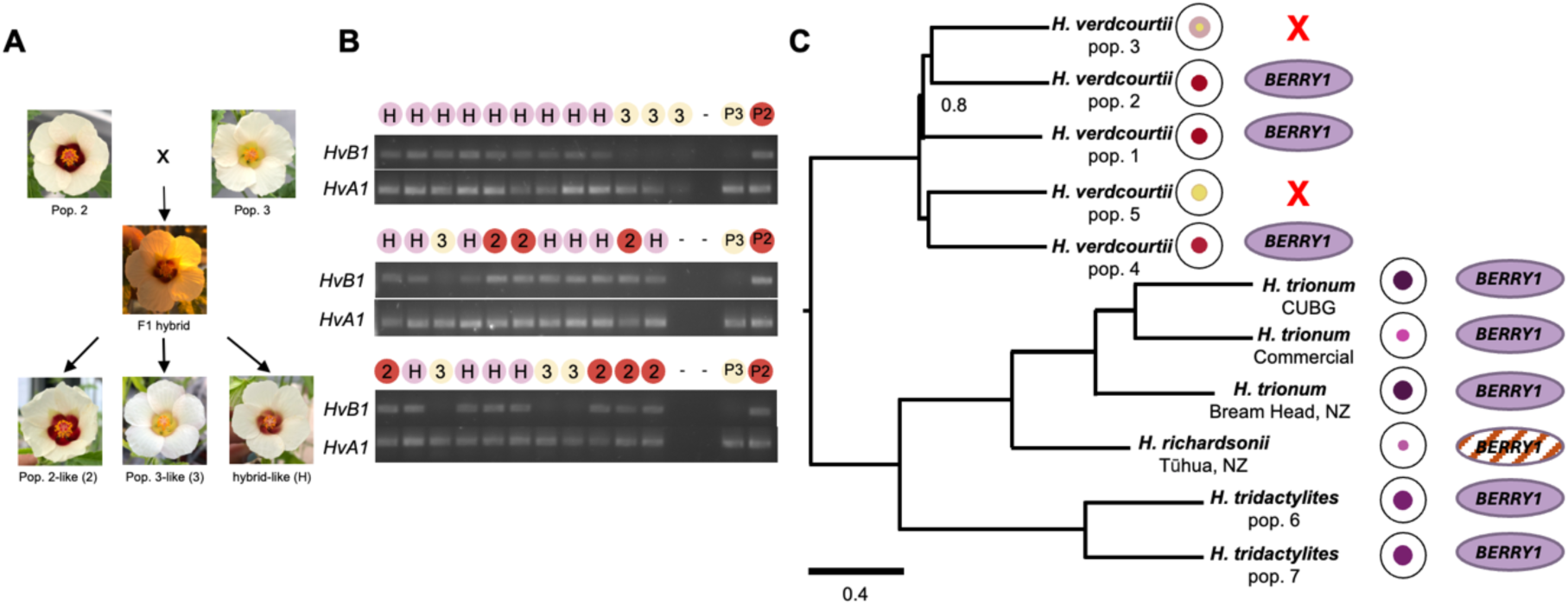
Presence of *HvBERRY1* correlates with pigmented bullseyes in *H. verdcourtii*. **(A)** F1 and F2 flower phenotypes of *H. verdcourtii* pop. 2 crossed *with H. verdcourtii* pop. 3. The F1 hybrid produced a flower with a pale pigmented bullseye. The F2 selfing population yielded three flower phenotypes that recapitulated the two parental and hybrid flower morphologies. **(B)** Genotyping of the *H. verdcourtii* pop. 2 x *H. verdcourtii* pop. 3 F2 population. Blind genotyping of 34 F2 individuals using *HvBERRY1*-specific primers revealed that the presence of *HvBERRY1* segregates with a pigmented bullseye. *HvB1 = H. verdcourtii BERRY1, HvA1 = H. verdcourtii ACTIN1*. **(C)** Multiple independent losses of the pigmented bullseye in the *Hibiscus trionum* Complex are a result of modifications to the *BERRY1* locus. In the *H. verdcourtii* clade, loss of *HvBERRY1* occurred twice and in both instances is correlated with a flower phenotype lacking a pigmented bullseye. In a third occurrence, a pigmented bullseye is absent in *H. richardsonii. HrBERRY1* transcripts are detected however several mutations within the coding region likely render it nonfunctional. Numbers next to nodes indicate support values. Scale bar represents number of changes per site.

### Bumblebee preference assays favor larger pigmented bullseyes

The fact that sister species but also distinct populations from the same species exhibit pigmented bullseyes of contrasting sizes raises the interesting possibility that a change in pattern dimension could influence pollinator selection, reduce genetic exchanges between populations and thus play a role in reproductive isolation and speciation. We previously showed that buff-tailed bumblebees (*Bombus terrestris* v. audax) could discriminate between 3D-printed plastic discs mimicking the proportions of *H. trionum* and *H. richardsonii* bullseye. The bumblebees had an innate preference for the discs with the larger bullseye, visiting those targets three out of four times when given the choice between the two types of patterns (Riglet *et al*. 2024). Real flowers are complex objects which can combine differences in shape, scent and temperature in addition to differences in pigmentation and those other features could influence pollinators’ response. To test bumblebees’ behavior in a more realistic context, we conducted preference tests experiments using real flowers from each species. We found that *B. terrestris* chose to land on *H. trionum* flowers over *H. richardsonii* ones 85% of the time (n= 40; 34 bees selected *H. trionum* vs. six bees selected *H. richardsonii*, p = 3.51 x 10^-7^, one-sample t-test) indicating that the preference for the larger bullseye was maintained or even reinforced in a real-flower context. This result demonstrates how a change at a single locus, *BERRY1*, can trigger a striking variation in pattern dimension and significantly influence pollinator behavior. The geographical distribution of *H. trionum* and *H. richardsonii* overlap in the North Island of New Zealand (Craven *et al*. 2011) and we have shown the two species are cross-fertile when hand pollinated in greenhouse conditions (Fig. 3A). In the future, it will be important to determine whether hybridization mediated by pollinating insects can also occur in the wild or whether the difference in bullseye size, combined with other factors, favors reproductive isolation.

## Conclusion

The occurrence of forms closely resembling each other across the tree of life supports the notion that similar features repeatedly arose independently through evolution. However, the molecular basis supporting phenotypical convergence is often unclear. Here, we used reduction in petal pattern dimensions across a small group of *Hibiscus*, the Trionum Complex, to unravel the molecular underpinnings of reduction in bullseye pigmentation. Relationships within the *Hibiscus* genus are well-known to be difficult to resolve (Pfeil & Crisp, 2005) but here we showed how the use of recent phylogenomic approaches can help illuminate the evolutionary history of previously recalcitrant groups, such as the Trionum Complex.

We found that the size of the pigmented area shrunk several times independently across the group. In at least three cases, this reduction was due to distinct changes affecting the same MYB-encoding locus, *BERRY1*, triggering either reduction in expression, premature stop codons of *HrBERRY1* in *H. richardsonii,* or complete gene loss of *HvBERRY1* from the genome of two populations of *H. verdcourtii*. This is consistent with the idea that regulators of the flavonoid pathway evolve faster than the structural genes they code for (Wheeler *et al*. 2022). This result also demonstrates how a change at a single locus, *BERRY1*, can trigger a striking variation in pattern dimension and significantly influence pollinator behavior as our data revealed that bumblebees exhibit a strong innate preference for the larger bullseye of *H. trionum* over the reduced one of its sister species. Whether repetitive reductions in bullseye size also impact the flower’s ability to deal with abiotic stresses, such as exposure to high levels of UV and dry environments, as observed in other species (Koski *et al*. 2015; Todesco *et al*. 2022), and whether it played a role in speciation events and diversification of the Trionum Complex remains to be established. Further investigations within the natural habitat of these *Hibiscus* species will provide valuable information such as the identification of species-specific pollinators, if any, and the frequency of self-fertilization events associated with possible low pollinator availability.

We also found that production of a pigmented bullseye in the Trionum Complex rests upon spatial restriction of MYB transcription factor activity. To date, all studies examining pigment pattern production in flowers found that the formation of colorful motifs relies on the precise control of *MYB* or *bHLH* expression (Albert *et al*. 2014) but spatial restriction of MYB transcription seems more common. In *M. lewisii,* a two-component activator-inhibitor system involving two MYB TFs, NEGAN and RTO, accounts for the production of pigmentation spots on *Mimulus* corollas (Ding *et al*. 2020). Consistently, in *Antirrhinum,* Petunia and maize, the R2R3 MYBs, rather than their bHLH partners, are often the main drivers of localized anthocyanin synthesis (Piazza *et al*. 2002; Schwinn *et al*. 2006; Albert *et al*. 2011). Understanding why MYBs rather than bHLHs tend to limit the production of pigment to subregions of the epidermis is unclear. It could reflect fundamental differences in the transcriptional regulation of both families, or instead indicate that the MYB components are less likely to have pleiotropic effects than their bHLH partner. Together, these findings underscore that evolution often recycles molecular mechanisms to generate petal diversity, revealing that the investigation of regulatory networks provide critical insights into how repeated evolutionary outcomes are achieved.

## Materials and Methods

### Plant material and growing conditions

Wildtype *H. trionum* L. CUBG seeds (voucher CGE00046422) were sourced from the Cambridge University Botanic Gardens, Cambridge, UK. *H. trionum* seeds from Bream Head, New Zealand (vouchers AK253689 & CGE00046417) were sourced from Dr. Brian G. Murray (University of Auckland). *H. trionum* Commercial seeds were supplied by Doekele G. Stavenga (Netherlands). Seeds of *H. richardsonii* (vouchers AK251841 & CGE00046420) were also sourced from Dr. Brian G. Murray (University of Auckland). Seeds of different populations of *H. verdcourtii* [population 1 (voucher CGE00046415), population 2 (voucher CGE00046414), population 3 (voucher CGE00080882), population 4 (voucher CGE00046413), population 5 (voucher CGE00046366)] and *H. tridactylites* [population 6 (voucher CGE00046364) and population 7 (voucher CGE00046363)] were given by Stephen B. Johnson (New South Wales, Department of Primary Industries). Seeds were soaked in 90°C H2O for 10 min and germinated in the dark at 30°C for 48 h. Seedlings were transferred to Levington High Nutrient M3 compost and grown under glasshouse conditions consisting of a 16 h light/8 h dark photoperiod at 25°C with a minimum radiance of 88 W.m^-2^. Wildtype species and transgenic lines used in this study are summarized in Table S1 and S2, respectively.

### Phenotyping

Open flowers (stage 5, Moyroud *et al*. 2022) were collected in the morning and imaged on black velvet using a Panasonic DMC-FZ38 camera (Panasonic Corporation). Dissected petals were imaged with a Keyence microscope (VHX5000 and VHX7000 models) fitted with a VH-Z100R lens. Petals with the adaxial side facing upward were scanned with a black and white background using the Epson Perfection V850 Pro scanner. Total petal and purple pigmented areas were measured from scanned images using ImageJ version 1.53 (https://imagej.net/ij/). For each flower, 3 petals were measured and a minimum of 10 flowers were measured per genotype.

### Flavonoid extraction and quantification

*Sample collection and extraction:* Stage 5 petals were dissected into proximal, boundary and distal regions and flash frozen, along with ovary tissue, in liquid nitrogen. For each sample, ∼40 mg of tissue was homogenized in a tissue lyser. 1 ml of acidic methanol (70:30 MeOH:1% acetic acid) was added to the tissue, vortexed and then incubated overnight on a rotating wheel at 4°C. Petal debris was pelleted for 3 min at 14680 RPM and the supernatant was collected and stored at 4°C until the following day. The extraction was repeated with 1 ml of acidic methanol (90:10 MeOH:1% acetic acid). Supernatants from both extractions were pooled before reducing to ∼250 μl by evaporation in a miVac centrifugal concentrator (Genevac). The final volume was measured before measurements were taken.

*Sample quantification:* Flavonols and anthocyanins were quantified using a spectrophotometer (SHIMADZU, UV-1800), with detection set between 349-350 nm and 529-530 nm, respectively. Three technical replicates (three dilutions) per extraction were measured. Concentrated extracts were diluted in 1 ml MeOH to measure flavonol concentration, and in 1 ml 90:10 MeOH:1N HCl to measure anthocyanin concentration. Absorbance units (AU) were converted to concentration (mg flavonol or anthocyanin equivalent per g of fresh weight tissue) based on the following calculation:

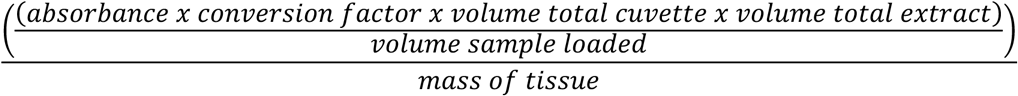

To determine conversion factors, standard curves for flavonol and anthocyanin equivalents were generated using the flavonol quercetin 3-D-galactoside (Cayman Chemicals item 18648) and anthocyanin cyanidin 3-O-glucoside (chloride) (Cayman Chemicals item 16406) as reference compounds.

### Crosses

For each cross, pollen was transferred by hand to the recipient flower. To avoid self-fertilization, recipient flowers without visible pollen on their stigma were selected for crossing.

1. *H. trionum x H. richardsonii*: five independent crosses of wildtype *H. trionum* and *H. richardsonii* were performed. F1 hybrids were identified based on the presence of a pigmented bullseye and microscopy imaging. F1 hybrid plants were allowed to self-pollinate and flower phenotypes of the next generation (F2) individuals were characterized.
2. *H. verdcourtii pop. 2 x H. verdcourtii pop. 3*: pollen was transferred by hand from a *H. verdcourtii* flower (St. George, red bullseye, “*H. ver2”*) to an open *H. verdcourtii* (St. George, pale yellow with pink halo bullseye, “*H. ver3”*). F1 hybrid plants were allowed to self-pollinate and flower phenotypes of the F2 generation were characterized.

### Genomic DNA extraction for whole genome sequencing

A total of 800 mg of mixed petal/bud tissue was divided into 50 mg aliquots for genomic DNA extraction. 50 mg aliquots of homogenized tissue were incubated with 1 ml CTAB buffer (2% w/v) + 10 µl 2-mercaptoethanol (Sigma M3148) at 55°C for 30 min. Supernatant was collected from pelleted samples and incubated with 10 mg.ml^-1^ RNase A (ThermoFisher Scientific EN0531) at 32°C for 30 min. DNA was extracted using 25:24:1 phenol-chloroform-isoamyl alcohol saturated with 10 mM Tris, pH 8.0, 1 mM EDTA (Sigma P0269) and 24:1 chloroform-isoamyl alcohol mixture (Sigma 25666). DNA was separated from protein and other cellular debris by carefully removing the upper aqueous layer after each extraction. The DNA was precipitated overnight at -20°C in chilled isopropanol and washed twice with 70% EtOH. DNA pellets were dried and resuspended in 30 µl low TE buffer (10 mM Tris HCl pH 8, 0.1 mM EDTA pH 8.0). Genomic DNA purity was analyzed using a NanoDrop (ThermoFisher Scientific) and Genomic DNA ScreenTape (Agilent 5067-3565 and 5067-3566) on the Agilent TapeStation 2200 System. For genome sequencing, a total of 162 µg HMW gDNA (>60,000 bp) was extracted at a concentration of 277.2 ng.µl^-1^ and a DIN value of 9.4. The sample was shipped on dry ice to Azenta Life Sciences, NGS Services Hope End, UK (formerly GeneWiz) for whole genome sequencing.

### Rapid genomic DNA extraction to isolate *HrBERRY1*

Fresh leaf tissue (∼1 cm^2^) was ground with a plastic pestle before addition of 250 µl Tris HCl-EDTA extraction buffer with 10% SDS. The samples were briefly vortexed and spun for 1 min at 14680 RPM to collect genomic DNA from the supernatant. DNA was precipitated with isopropanol and the pellet was washed with 70% EtOH. The pellet was dried before resuspension in 40 µl ddH2O.

### Genotyping the F2 progeny from wildtype *H. trionum* and *H. richardsonii* crossing

A single bud per individual was collected from wildtype *H. trionum* and *H. richardsonii* parents, and from 27 F2 individuals randomly chosen among >100 F2 individuals that had been phenotyped and each assigned to one of three phenotypes (*H. trionum*-like, *H. richardsonii*-like, hybrid-like). Genomic DNA extraction from selected individuals was performed following the CTAB protocol used for whole genome sequencing. The presence of *HtBERRY1* or *HrBERRY1* gene copies were determined by PCR using allele-specific primers (Table S3) for *HtBERRY1* or *HrBERRY1*. A 1200 bp band indicated the presence of the *HtBERRY1* allele and a 400 bp band indicated the presence of the *HrBERRY1* allele. Samples were genotyped in a blind manner (phenotyped by MTSY and genotyped by AMF who did not know the phenotypes). To confirm the genotyping results, a further 11 F2 individuals with a *H. richardsonii*-like phenotype were tested. *HtACTIN1* was included as a positive control for the presence of genomic DNA in all samples tested.

### Genotyping the F2 progeny from *H. ver2* and *H. ver3* crossing

Fresh leaf tissue (∼1 cm^2^) was collected from *H. ver2* and *H. ver3* parents, and 33 F2 individuals. The tissue was ground with a plastic pestle before addition of 250 µl Tris HCl-EDTA extraction buffer with 10% SDS. The samples were briefly vortexed and spun for 1 min at 14680 RPM to collect the genomic DNA from the supernatant. DNA was precipitated with isopropanol and the pellet was washed with 70% EtOH. The pellet was dried before resuspension in 40 µl ddH2O. Samples were genotyped in a blind manner to check for the presence of *HvBERRY1*. The presence of *HvBERRY1* was determined by PCR using *HvBERRY1*-specific primers (Table S3). F2 samples with a pigmented red bullseye produced a 421 bp band that indicated the presence of *HvBERRY1* while those without a pigmented bullseye did not produce a PCR product. *HvACTIN1* was included as a positive control for the presence of genomic DNA in all samples tested.

### Tissue collection, RNA extraction and Illumina sequencing to produce the mixed tissue transcriptome for phylogenetic analyses

Whole bud, flowers and fruit tissue were collected from each of the following species and populations: wildtype *H. trionum* (voucher CGE00046422), *H. trionum* diploid New Zealand naturalized race (voucher AK253689, CGE00046417), *H. trionum* Commercial Variety (voucher CGE00080883), *H. richardsonii* (voucher AK251841, CGE00046420), *H. verdcourtii* (vouchers CGE00046415, CGE00046414, CGE00080882, CGE00046413 and CGE00046366), *H. tridactylites* (vouchers CGE00046364 and CGE00046363). Upon collection, bud tissue was flash frozen in liquid nitrogen. Per species/population, three biological replicates were collected with each biological replicate corresponding to one individual plant. Frozen samples were homogenized using a pestle and mortar and total RNA was extracted using the Spectrum Plant Total RNA Kit (Sigma STRN250). ∼40 mg of tissue was used for each RNA extraction. An on-column DNase I treatment (Sigma DNASE70) was performed between the first two column washes. In the final step, RNA was eluted in 30 µl nuclease-free H2O (Promega MC1191). RNA purity was analyzed using a NanoDrop (ThermoFisher Scientific) and the RNA ScreenTape assay (Agilent 5067-5576, 5067-5577, 5067-5578) on the Agilent TapeStation 2200 system.

Samples with minimum 1.7 µg total RNA (>50 ng µl^-1^), and A260/280 value of 2.0 were shipped on dry ice to Azenta Life Sciences, Leipzig (formerly GeneWiz) for RNA-seq library preparation and sequencing. Library preparation was performed with polyA selection and sequencing using an Illumina HiSeq machine (2x 150 bp configuration, ∼350 M raw paired-end reads per lane, at least 30M reads per sample).

### Phylogenetic transcriptome analysis

Raw sequence reads for the outgroups *(Hibiscus cannabinus, Hibiscus syriacus* and *Hibiscus sabdariffa)*, were downloaded from the NCBI Short Read Archive (SRA) accessions (SRX1084555 [Zhang *et al*. 2016], SRX2142897 [Kim *et al*. 2017], SRX3341155 [Loo *et al*. 2016]).

The sequences were assembled using the same procedure as the *H. trionum* stage 1 petal transcriptome.

We reduced the redundancy for each transcriptome using cd-hit-est v.4.7 (Fu *et al*. 2012), with the settings “-c 0.99 -n 10 -r 0”. Initial homology inference was conducted by performing an all-by-all blastn v.2.7.1 with the settings “--e-value 10 –max_target_seqs 1000”. Markov clustering, as implemented in the program mcl (van Dongen, 2000) with an inflation value of 1.4, was used to refine the clusters further. We then performed two rounds of phylogeny-based data cleaning. In the first round, the sequences were aligned using MAFFT v.7.407 (Katoh & Standley, 2013) with the settings “—auto –maxiterate 1000” the alignments were cleaned for 10% column occupancy using the program pxclsq with the settings “-p 0.1” from the phyx package (Brown *et al*. 2017). A tree for each cleaned alignment was inferred using maximum likelihood as implemented in FastTree v.2.1.1 (Price *et al*. 2010) with the settings “-gtr -nt”. The resulting trees had any tips with an absolute branch length of 0.4 subs/site or a relative branch length of 0.6 subs/site removed. Only the transcript with the most informative sites was retained for any instance where the same taxa were sister to one another. Any subclades containing at least 4 taxa and were on a branch with a 1.0 subs/site length were divided into separate homolog trees. During the second round of cleaning, the same process was performed, however, the maximum likelihood tree inference was conducted using IQtree v.1.6.12 (Nguyen *et al*. 2015) using the best inferred model of evolution (Kalyaanamoorthy *et al*. 2017) and 1000 ultrafast bootstrap replicated. The orthologs were extracted from the trees using the Maximum Inclusion approach with an absolute branch length setting of 0.4 subs/site and a relative branch length setting of 0.6 subs/site. A coalescent based maximum quartet support species tree was estimated using Astral v.5.7.3.

### Tissue collection, RNA extraction and Illumina sequencing for *H. trionum* stage 1 petal transcriptome

Tissue was collected from five individual wildtype *H. trionum* plants to generate five biological replicates: for each biological replicate, stage 1 petals were harvested and dissected to separate proximal and distal region. Tissue from these 10 samples were flash frozen in liquid nitrogen, ground to a fine powder, and total RNA extracted using the Spectrum Plant Total RNA kit (Sigma STRN250) with 40 mg of starting material. An on-column DNase I (Sigma DNASE70) treatment was performed between the first two column washes. In the final step, RNA was eluted with 30 µl nuclease-free H2O (Promega MC1191). RNA purity was analyzed using a NanoDrop (ThermoFisher Scientific) and the RNA ScreenTape assay (Agilent 5067-5576, 5067-5577, 5067- 5578) on the Agilent TapeStation 2200 system. Samples with minimum 1.75 µg total RNA (>50 ng.µl^-1^), and A260/280 value of 2.0 were shipped on dry ice to Azenta Life Sciences, Leipzig (formerly GeneWiz) for RNA-seq library preparation and sequencing. Library preparation was performed with polyA selection and sequencing using an Illumina HiSeq machine (2x 150bp configuration, ∼350M raw paired-end reads per lane, at least 30M reads per sample).

### *H. trionum* stage 1 petal transcriptome assembly and differential gene expression analysis

RNA-seq reads were used to assess gene expression and compare transcript abundance between proximal and distal regions of stage 1 (S1) petal primordia, as defined in Moyroud *et al*. 2022. First, a reference transcriptome (Supplementray Data 2) was created by combining the raw reads from S1 proximal and distal petal tissue of a single plant. To produce this reference transcriptome the raw reads were cleaned, assembled into a database with chimeric sequences removed, and their coding regions for transcripts predicted using the previously excluded chimeric sequences. Cleaning raw reads used Rcorrector v1.0.3 (Song & Florea, 2015), to remove sequencing errors, and Trimmomatic v0.38 (Bolger *et al*. 2014), to filter for quality and remove the adapters added during library preparation. The cleaned reads were then assembled into a transcriptome with Trinity v2.8.3 (Grabherr *et al*. 2011). The procedure followed Yang and Smith (2013) with a database composed of the genomes of *Gossypium raymondii*, *Arabidopsis thaliana*, and *Theobroma cacao*. Chimeric sequences were filtered by BLAST^®^ v2.2.31 (Altschul *et al*. 1997) and isoforms and assembly artifacts removed using Corset v1.07 (Davidson & Oshlack, 2014). Finally, coding regions were predicted using the program Transdecoder v5.3.0 (Haas *et al*. 2013) and a BLAST database composed of the same taxa as the chimera database.

The reference transcriptome was then used as a guide for transcriptome assembly corresponding to each biological replicate. Raw reads from each sample were first mapped onto the reference transcriptome using Kallisto v.0.44.0 (Bray *et al*. 2016). The transcript level- abundance was converted into counts using the program tximport (Soneson *et al*. 2015). Fold change and dispersion of these counts were calculated using DESeq2 (Love *et al*. 2014). Finally, the top blastn hit between the transcriptome sequences and the NCBI nr database was used for automatic gene annotation. Genes with log2-fold change <-2 or >2 and adjusted p-value < 10^-5^ were considered differentially expressed. Results of the Differential Gene Expression analysis and assembled transcripts from stage 1 *H. trionum*, CUBG petals are provided in Supplementary Data 1 & 2.

### Phylogenetic analysis to establish the identity of selected R2R3 MYBs

The amino acid sequences of HtBERRY1 (HRI_002225000), HtBERRY2 (HRI_002225200) and HtCREAM1 (HRI_001263100) were used to conduct a blast search against the genome of *H. trionum*available on GenBank (https://www.ncbi.nlm.nih.gov/datasets/genome/GCA_030270665.1/). This yielded another six *HtBERRY*-like sequences (HRI_00224000, HRI_002224800, HRI_002224600, HRI_000181400, HRI_001579800, HRI_002131000) and another three *HtCREAM*-like sequences (HRI_003566500, HRI_00625700, HRI_001075200). Those 12 *H. trionum* sequences were combined with protein sequences of subgroup 6 MYB genes known to regulate anthocyanin production in other species: ROSEA1 (DQ275529), ROSEA2 (DQ275530) and VENOSA (DQ275531) from *Antirrhinum majus*; PELAN (KJ011144) and NEGAN (KJ011145) from *Mimulus lewisii*; MYB113-like (XM_012977147) from *Mimulus guttatus*; DEEP PURPLE (HQ116169) and PURPLE HAZE (HQ116170, HQ428100) from *Petunia hybrida* as well as amino acid sequences of all MYB genes identified in the genome of *Arabidopsis thaliana*, *Gossypium raimondii* and *Theobroma cacao* were acquired from Jiang *et al*. 2020, He *et al*. 2016 and Du *et al*. 2022, respectively. In total, 483 MYB sequences were used for phylogenetic reconstructions and those are provided in Supplementary Data 3. The 483 sequences were aligned using MAFFT v7.490 with the setting “—auto –maxiterate 1000”. The alignment was then cleaned for a minimum of 10% column occupancy using the program pxclsq from the phyx package. A maximum likelihood tree was then inferred from the cleaned alignments using the WAG model of evolution, with gamma rate variation and 1000 ultrafast bootstrap replicates as support (Hoang *et al*. 2018).

### cDNA synthesis

cDNA was synthesized from 0.5 ug total RNA using SuperScript III Reverse Transcriptase (Invitrogen). Oligo(dT)15 primers were used for first strand cDNA synthesis for gene isolation and measuring expression by RT-PCR or qPCR. First-strand cDNA synthesis followed the manufacturer’s guidelines, with incubation with SuperScript III RT at 50°C for 1 hr followed by an inactivation step at 70°C for 15 min.

### Construction of plant expression vectors

The coding sequences of *HtDFR1, HtBERRY1* and *HtCREAM1* were amplified by PCR using Q5 High-Fidelity DNA polymerase with primers listed in Table S3. Each PCR product was cloned into the EcoRV-digested pBluescript KS(-) vector to produce an intermediate vector. Restriction enzyme digestion of the intermediate vector was used to extract the coding sequence of each gene and insert it into a modified pGREENII (for *HtDFR1* and *HtCREAM1* overexpression) or pCAMBIA (for *HtBERRY1* overexpression) containing a 2x35S CaMV promoter driving transgene expression. The plant expression vectors also express a fluorescent eYFP under control of the promoter region of *AtUBIQUITIN10* (PromUBQ10). Final vectors were sent to Azenta (formerly Genewiz) for Sanger sequencing to validate the final construct prior to plant transformation. The final vectors used for plant transformation are summarized in Table S4.

### Plant transformation

Electrocompetent *Agrobacterium tumefaciens* LBA4404 cells were transformed with plant expression vectors pVT9 *(2x35S::HtDFR1)*, pAF29 *(2x35S::HtCREAM1)* or pAF52 *(2x35S::HtBERRY1)*. Transformed cells were selected on LB agar plates containing 50 mg.l^-1^ kanamycin and 25 mg.l^-1^ streptomycin after 48 hours incubation in the dark at 30°C. Transgenic *H. trionum* lines were produced using the protocol described in Moyroud *et al*. 2022 with the following modification: 100 mg.l^-1^ of carbenicillin was used instead of 250 mg.l^-1^ of cefotaxime for the MS Hib plates.

### Gene expression analysis using quantitative RT-PCR

To quantify gene expression levels during petal development in wildtype *H. trionum* and *H. richardsonii*, petals from stages 1 to 4 were dissected into proximal and distal regions. Stage 3 petals were dissected into proximal and distal regions to measure expression in the transgenic lines of *35S::HtDFR1* OE and KD*, 35S::HtCREAM1* and *35S::HtBERRY1*. Three biological replicates were collected from individual plants for each stage. Frozen petal tissue was ground to a fine powder and total RNA was extracted using the Spectrum Plant Total RNA kit (Sigma STRN250) with ∼40 mg of ground tissue used as starting material for each RNA extraction. cDNA was synthesized from total RNA using SuperScript III Reverse Transcriptase (Invitrogen) following the manufacturer’s guidelines. Quantitative real-time PCR was performed using a Roche LightCycler 480 machine and the Luna Universal qPCR Master Mix (New England BioLabs). *HtACTIN1* was used as a housekeeping gene for all biological replicates and stages as in Moyroud et al. 2022. Data was analyzed using a modified delta Ct method, considering the real efficiency of each primer pair and one reference gene for normalization (Pfaffl, 2001; Ganger *et al*. 2017). Statistical analyses were calculated in RStudio (Version 1.1.1717). A Student’s t-test, Welch’s t-test or Wilcox’s t-test was used depending on normality and homogeneity of variance of the samples.

### Bumblebee behavior assay

Experiments with flower-naive buff-tailed bumblebees (*Bombus terrestris* v. audax; Research Hive, BioBest UK) were conducted using the design published in Moyroud *et al*. 2017. Each colony was fed daily with fresh 15% sucrose solution, and twice a week with pollen grains (The Happy Health Company). Foragers were hand-marked with water-based Thorne queen marking paints in various colors to distinguish them during the experiments. For the preference tests, one wildtype *H. trionum* and one *H. richardsonii* open flower were each placed in a tube containing water equidistant from the hive entrance. Per test, one naïve bumblebee was released from the hive into the arena and the flower it chose to land on first (preference) was recorded. Forty individual bumblebees were tested for their flower preference. Statistical differences were calculated using a one sample t-test [RStudio (Version 1.1.1717)].

## Acknowledgments

We thank Brian. G. Murray (University of Auckland) and Stephen B. Johnson (NSW Department of Primary Industries) for the generous gift of *H. richardsonii*, *H. trionum* Bream Head, *H. tridactylites* and *H. verdcourtii* seeds from Australia and New Zealand. We thank Doekele G. Stavenga for *H. trionum* Commercial seeds. We also thank all members of the Moyroud group for regular discussions on the results presented here, particularly Erin Doody, Edwige Berthelot, Elena Salvi, and Philip Carella (JIC) for useful feedback and comments on earlier versions of the manuscript. We acknowledge support from the entire SLCU community including general lab support, horticulture, and facilities teams.

## Author Contributions

E.M. conceptualized and designed the project. M.T.S.Y., A.L.M.F., V.T., S.Z. and L.R. performed experiments and data analysis. M.T.S.Y. validated experiments and performed statistical analysis. J.F.W. analyzed the transcriptome data. M.T.S.Y. and E.M. prepared figures and wrote the manuscript, with input from all authors.

## Data, Code and Material Availability

The coding and genomic sequences of all *H. trionum* genes mentioned in this study can be accessed via the genome of *H. trionum* available on GenBank (https://www.ncbi.nlm.nih.gov/datasets/genome/GCA_030270665.1/) using the HRI_ reference numbers provided in the Material & Methods sections. The coding sequences of HrDFR1, HrFLS2 and HvBERRY1 have been deposited in GenBank under the accession number PQ450317, PQ450318 and PQ450319, respectively. Transcriptomic data associated with the phylogenomic analysis of the Trionum complex and the comparative analysis of gene expression between proximal and distal petal regions of *H. trionum* stage 1 petal have been deposited on GenBank SRA (https://www.ncbi.nlm.nih.gov/sra). The amino acid sequences used for assessing the placement of HtBERRY1, HtBERRY2 and HtCREAM1 in the MYB family phylogeny are provided in Supplementary Data 3. The algorithms and codes used for phylogenomic analyses, RNAseq data analysis and differential gene expression analysis have been published elsewhere, as indicated in the references provided in the Material & Methods section.

