## Supplementary Figures and Tables for "The genetic basis of replicated bullseye pattern reduction across the Trionum Complex"

**Supplementary Data 1**. Differential gene expression analysis between proximal and distal regions of stage 1 *H. trionum* petal primordia. Data related to Fig S4.

**Supplementary Data 2**. Assembled transcripts from *H. trionum*, stage 1 petals. Data related to Fig S4.

**Supplementary Data 3**. Amino acid sequences of the 483 MYB sequences used to assess the phylogenetic placement of HtBERRY1, HtBERRY2 and HtCREAM1. Data related to Fig S5.

**Supplementary Data 4**. Complete MYB family tree generated using the 483 sequences from Supplementary data 3. Data related to Fig S5.

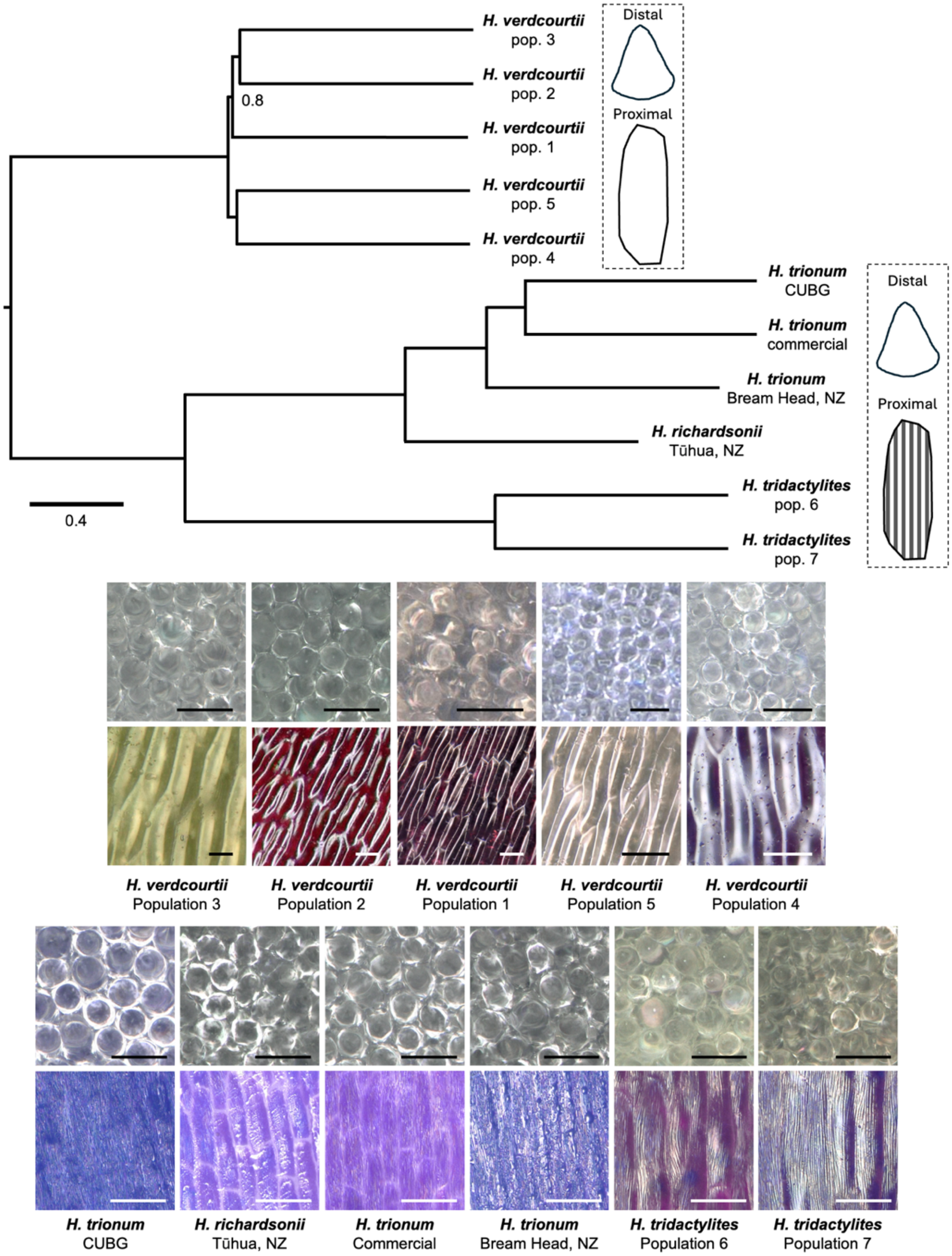


**Fig. S1.** Cell shape and texture across the adaxial petal epidermis of the 11 different accessions from the Trionum Complex used in this study (see Fig. 1). Unless stated, all nodes are perfectly supported. Branch lengths are shown in coalescent units. All accessions display tabular cells in the proximal petal region and smooth conical cells in the distal region. The proximal tabular cells in the proximal region are covered with a striated cuticle, except for the 5 *Hibiscus verdcourtii* accessions that display smooth tabular cells in the proximal petal region. Scale bar = 50uM.

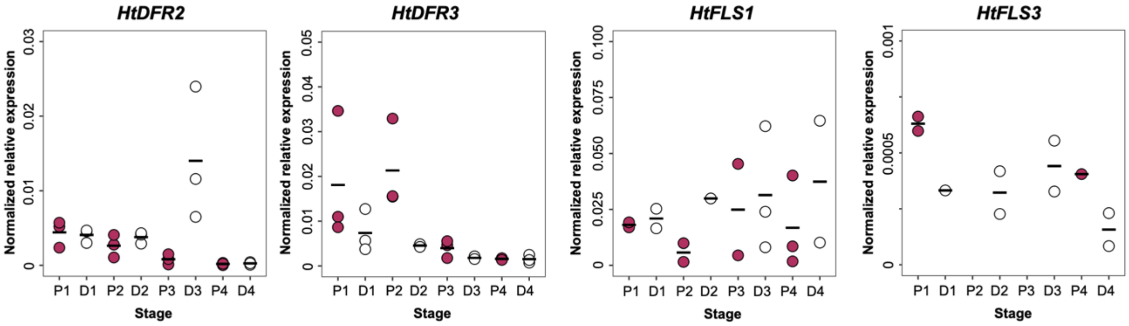


**Fig. S2.** qRT-PCR analysis of the expression of *HtDFR2, HtDFR3, HtFLS1 and HtFLS2* throughout bud development in proximal (P1 to P4, purple dots) and distal (D1 to D4, white dots) petal tissue of wildtype *H. trionum*. Three biological replicates were extracted per stage and each data point indicates an average of three technical replicates, horizontal bars indicate mean relative expression values.

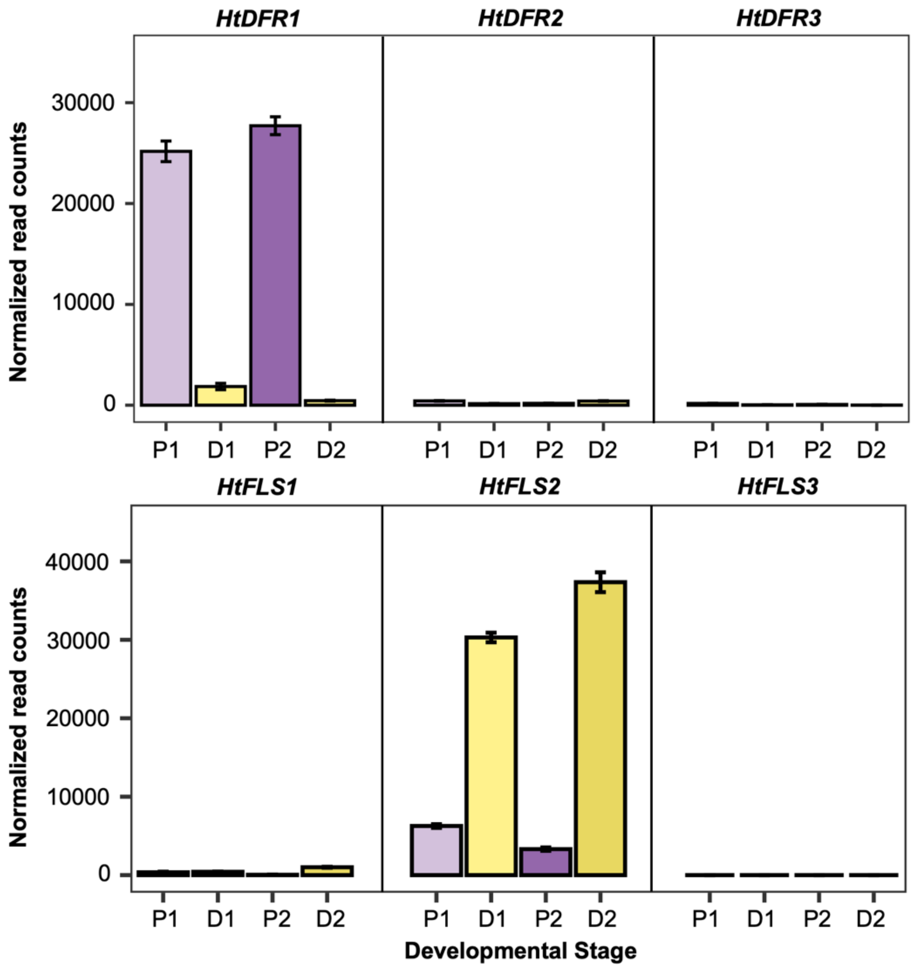


**Fig. S3.** Normalized read counts for 3 *HtDFR* homologs and 3 *HtFLS* homologs in proximal (P1 and P2) and distal (D1 and D2) petal tissue of wildtype *H. trionum* at stage 1 and stage 2 (before and after bullseye pigmentation emergence, respectively). The y-axis shows the normalized read counts in Transcripts Per Million (TPM).

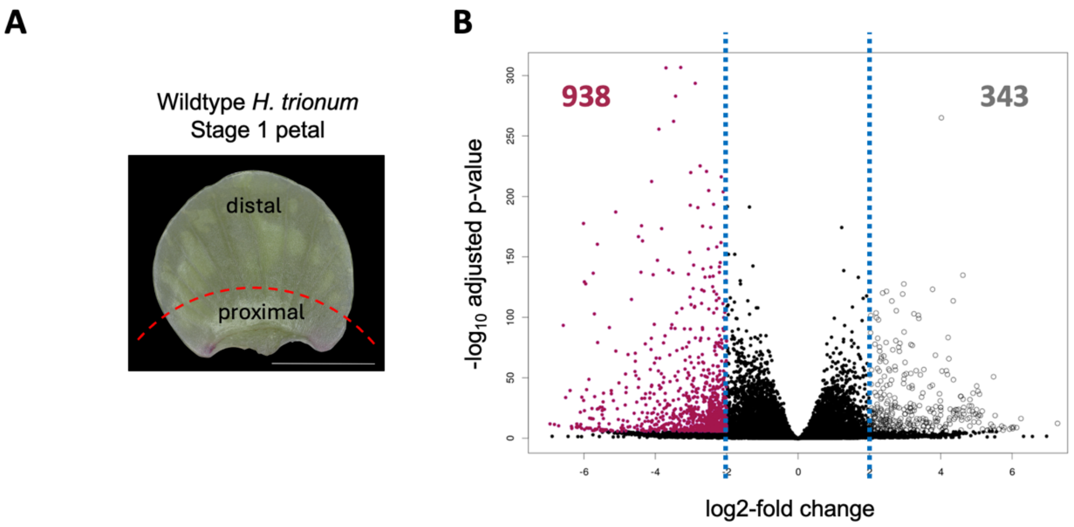


**Fig. S4.** Identification of differentially expressed genes between the proximal and distal regions of stage 1 petal primordia in wildtype *H. trionum* **(A)** Picture of a dissected wildtype *H. trionum* stage 1 petal showing the two regions used for RNA extraction and transcriptome analysis: region above the red dotted line = distal region, region below the red dotted line = proximal region. **(B)** Volcano plot displaying genes identified in the transcriptome analysis of stage 1 petals. Genes preferentially expressed in the proximal region (>4x difference compared to distal region) are represented by purple dots while genes preferentially expressed in the distal region (>4x difference compared to proximal region) are represented by white dots. The blue lines represent the log2-fold thresholds used for differential expression. Genes considered not differentially expressed between proximal and distal regions are depicted with black dots.

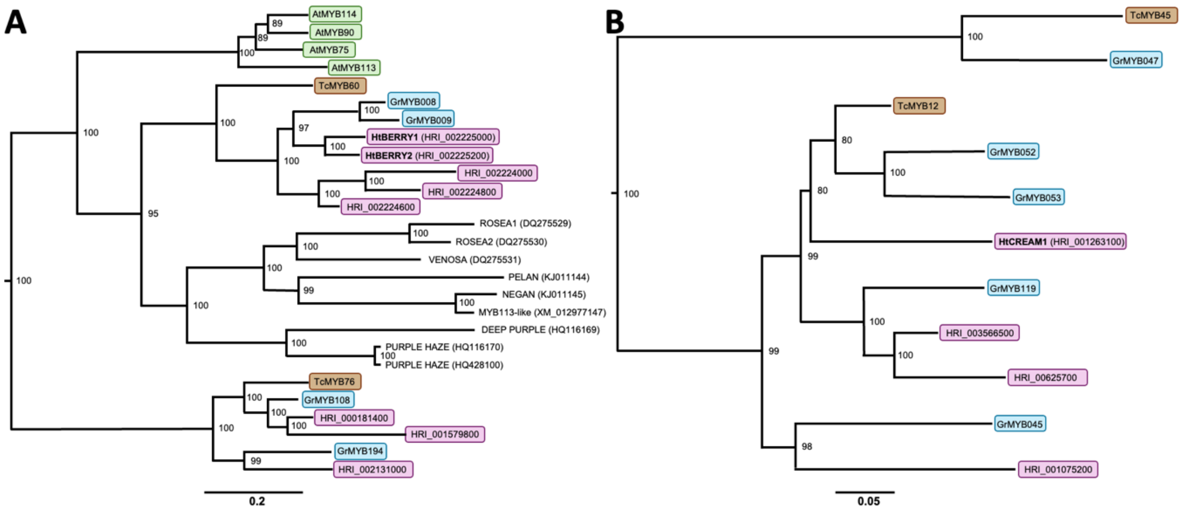


**Fig. S5.** Placement of *HtBERRY1*, *HtBERRY2* and *HtCREAM1* within the MYB family phylogenetic tree. **(A)** Phylogenetic tree depicting the position of *HtBERRY1 and HtBERRY2* within the subgroup 6 R2R3-MYB cluster**. (B)** Phylogenetic tree depicting homology relationships between *HtCREAM1* and its three *H. trionum* paralogs and R2R3-MYBs from cotton and cocoa. *H. trionum* genes are highlighted in purple, cotton (*Gossypium raimondii*) genes are highlighted in blue, cocoa (*Theobroma cacao*) genes are highlighted in brown and *Arabidopsis thaliana* genes are boxed in green. Ultrafast Bootstraps values are given next to each node. Scale bar represents number of changes per site. Subgroup 6 R2R3-MYBs from other species, known to regulate anthocyanin production in the flowers of other species have also been included in (A). *Antirrhinum majus*: ROSEA1, ROSEA2, VENOSA; *Mimulus lewisii*: PELAN, NEGAN; *Mimulus guttatus*: MYB113-like; *Petunia hybrida*: DEEP PURPLE, PURPLE HAZE. The trees presented in A and B are part of a larger R2R3-MYB phylogenetic analysis we conducted, including all known R2R3-MYBs from *A. thaliana*, *G. raimondii* and *T. cacao*. The sequences used for this analysis are provided a Supplementary Data 3 and the complete phylogeny is provided as Supplementary Data 4.

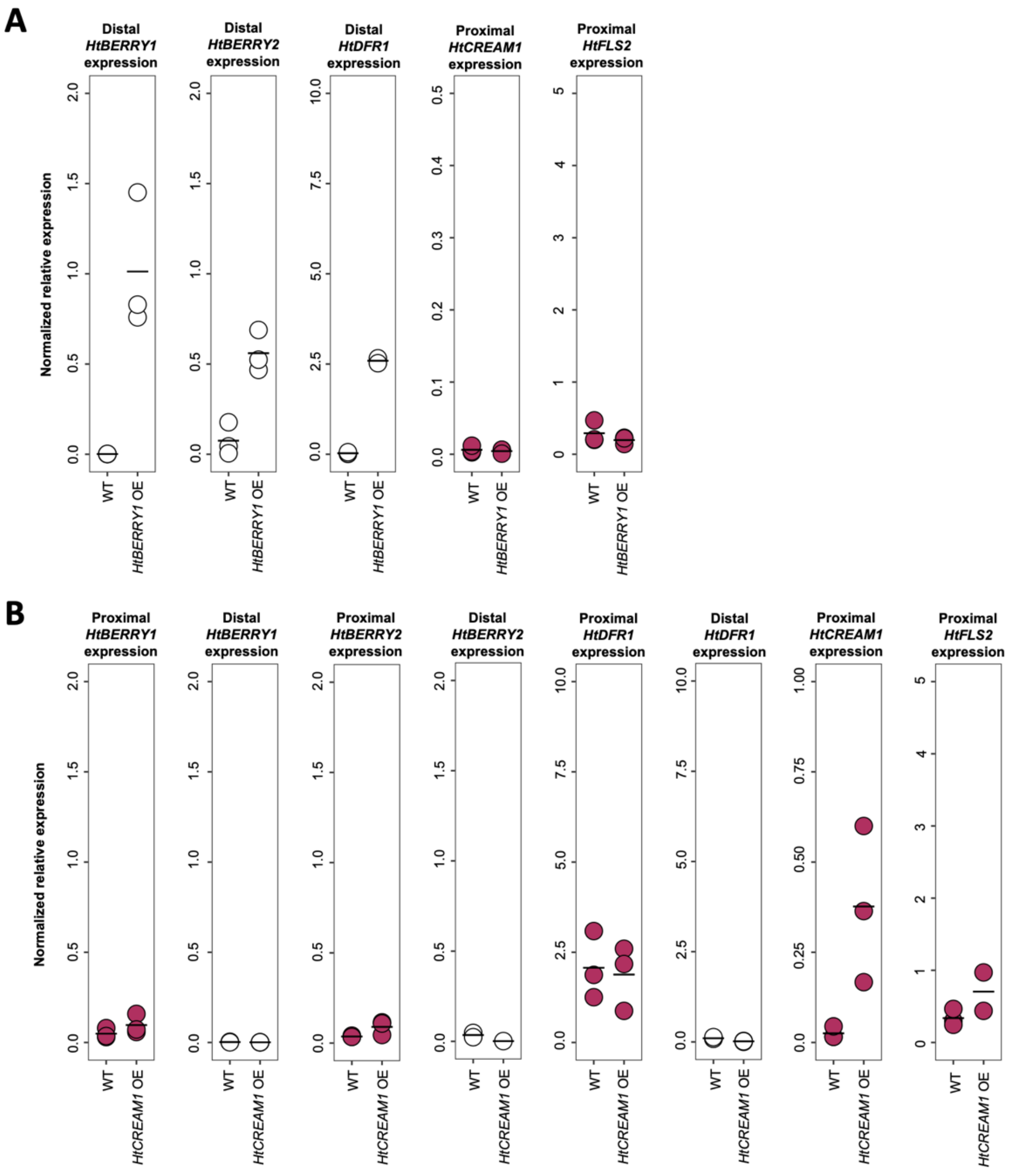


**Fig. S6.** qRT-PCR analysis of the expression of flavonoid-related structural and MYB transcription factors in distal (white dots) or proximal (purple dots) regions of wildtype and **(A)** *HtBERRY1* or **(B)** *HtCREAM1* OE *H. trionum* lines. Three biological replicates were extracted per stage and each data point indicates an average of three technical replicates, horizontal bars indicate mean relative expression values.

| Species | Description | Provenance | Voucher |
|---|---|---|---|
| *H. trionum* | Wildtype | Cambridge University Botanic Garden collection | CGE00046422 |
| *H. trionum* | Diploid New Zealand naturalized race | Bream Head, New Zealand | AK253689, CGE00046417 |
| *H. trionum* | Commercial | Netherlands | CGE00080883 |
| *H. richardsonii* | Wildtype | Mayor Island (Tuhua), New Zealand | AK251841, CGE00046420 |
| *H. verdcourtii* | Population 1 | Emerald, Queensland | CGE00046415 |
| *H. verdcourtii* | Population 2 | St. George, Queensland | CGE00046414 |
| *H. verdcourtii* | Population 3 | St. George, Queensland | CGE00080882 |
| *H. verdcourtii* | Population 4 | Theodore, Queensland | CGE00046413 |
| *H. verdcourtii* | Population 5 | Narrabri, NSW | CGE00046366 |
| *H. tridactylites* | Population 6 | Narrabri, NSW | CGE00046364 |
| *H. tridactylites* | Population 7 | Jimbour, Queensland | CGE00046363 |

**Table S1.** *Hibiscus* species used in this study.

| Line | Plant Background | Construct | Description |
|---|---|---|---|
| AF29.1 | *H. trionum* | pAF29 | *HtCREAM1* overexpression |
| AF52.1 | *H. trionum* | pAF52 | *HtBERRY1* overexpression |
| VT9.17 | *H. trionum* | pVT9 | *HtDFR1* overexpression |
| VT9.6 | *H. trionum* | pVT9 | *HtDFR1* overexpression |

**Table S2**. *H. trionum* transgenic lines generated in this study.

| Gene | Sequence | Purpose |
|---|---|---|
| HtDFR1-qF | TGCCGCCTAGCTTGATTACC | qPCR |
| HtDFR1-qR | CACGAACTGCCCTTGCCTAATA | qPCR |
| HtDFR2-qF | ACCTCCTTTAGTGGTTGGTCC | qPCR |
| HtDFR2-qR | TCCCAGTGATGGGAGAAAGTG | qPCR |
| HtDFR3-qF | TGGGACTTCTGCAACGAGAATAA | qPCR |
| HtDFR3-qR | ATCGGAGGCAGTGGAACATAAA | qPCR |
| HtFLS1-qF | GGAAGAGAAAGAGGTGTACGCTAAG | qPCR |
| HtFLS1-qR | CAGCCCAGTTCTTCTTCCCATTA | qPCR |
| HtFLS2-qF | GAATACGCTAAGCACATGCATGG | qPCR |
| HtFLS2-qR | CATGTCGTCCCCACCTAGAG | qPCR |
| HtFLS3-qF | GCGCCCTCGTTGTTCATATTG | qPCR |
| HtFLS3-qR | TCCGAGTTCTCTCCTTGTCTAC | qPCR |
| HtBERRY1-qF | GGTAGACTGCCTGGAAGAACAG | qPCR |
| HtBERRY1-qR | GATGAGGTTTCGGGTTCGAGTTA | qPCR |
| HtBERRY2-qF | GGTAGACTGCCAGGAAGAACAT | qPCR |
| HtBERRY2-qR | TGGATGAGGTTTCGGGTTTGA | qPCR |
| HtCREAM1-qF | GGACCGGGTTTCTTTCTGCTTAT | qPCR |
| HtCREAM1-qR | GTCATCTCCACTTGCTGGTACAC | qPCR |
| HrBERRY1-qF | GGTAGACTGCCTGGAAGAACAG | qPCR |
| HrBERRY1-qR | GATGAGGTTTCGGGTTCGAGTTA | qPCR |
| HrBERRY2-qF | GGTAGACTGCCAGGAAGAACAT | qPCR |
| HrBERRY2-qR | TGGATGAGGTTTCGGGTTTGA | qPCR |
| HrCREAM1-qF | GGACCGGGTTTCTTTCTGCTTAT | qPCR |
| HrCREAM1-qR | GTCATCTCCACTTGCTGGTACAC | qPCR |
| HtBERRY1-F | TTTGAGGGTTAGTGTTAAACAGCTACG | genotyping |
| HtBERRY1-R | AAACTTCTGGCTATAAGTTGAACACATTC | genotyping |
| HrBERRY1-F | CCTCGGTAACAGGTAACTCATTAG | genotyping |
| HrBERRY1-R | GGTCGAGTATGGTCCAATGG | genotyping |
| HvBERRY1-F | GATCAATCCTTCCGACAGTAACTCAAACC | genotyping |
| HvBERRY1-R | CTATAGGTTGAACACATTCCAGAACTCTGC | genotyping |
| HvACTIN1-F | CCCAGATCATGTTTGAGACCTT | genotyping |
| HvACTIN1-R | ACCGGAATCCAGCACAATAC | genotyping |

**Table S3.** Primer sequences used in this study.

| Vector | Recombinant DNA | Reference |
|---|---|---|
| pSG55 | Modified pCAMBIA1300 with 2xp35S and pUBQ10::eYFPmyr | This study |
| pAF29 | 2xp35SS::HtCREAM1 in pSG55 | This study |
| pAF52 | 2xp35SS::HtBERRY1 in pSG55 | This study |
| pVT9 | 2xp35SS::HtDFR1 in pSG55 | This study |

**Table S4**. Plant expression vectors generated for this study.
