## Supplementary figures and images for "The genetic basis of replicated bullseye pattern reduction across the Trionum Complex"

### Supplementary Data 4

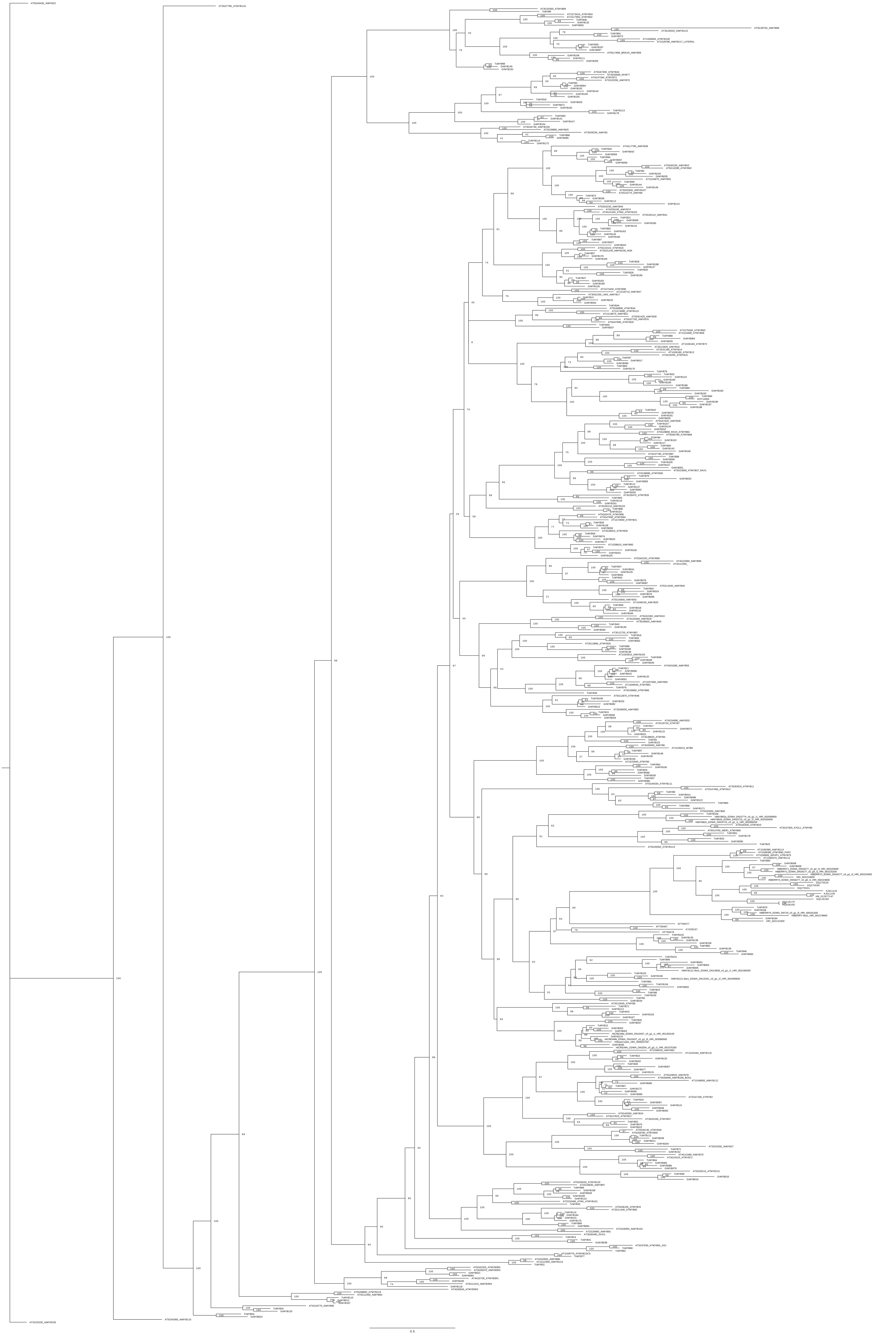
